# Benchmarking Alignment Strategies for Hi-C Reads in Metagenomic Hi-C Data

**DOI:** 10.1101/2025.07.30.667754

**Authors:** Yuqiu Wang, Wenxuan Zuo, Jiawei Huang, Fengzhu Sun, Yuxuan Du

## Abstract

**Background:** Metagenomics combined with High-throughput Chromosome Conformation Capture (Hi-C) offers a powerful approach to study microbial communities by linking genomic content with spatial interactions. Hi-C enhances shotgun sequencing by revealing taxonomic composition, functional interactions, and genomic organization from a single sample. However, aligning Hi-C reads to metagenomic contigs presents challenges, including the varying insert size of Hi-C paired-end reads, multi-species complexity, and gaps in assemblies. Although many benchmark studies have evaluated general alignment tools and Hi-C data alignment, none have specifically addressed metagenomics Hi-C data.

**Results:** Here, we selected seven alignment strategies that have been used in Hi-C analyses: BWA MEM -5SP, BWA MEM default, BWA aln default, Bowtie2 default, Bowtie2 –very-sensitive-local, Minimap2 default, and Chromap Hi-C default. We benchmarked them on one synthetic and seven real-world environments, and evaluated these tools based on the number of inter-contig Hi-C read pairs and their influence on downstream tasks, such as binning quality.

**Conclusion:** Our findings show that BWA MEM -5SP consistently outperforms other tools across all environments in terms of inter-contig read pairs and binning quality, followed by BWA MEM default. Chromap and Minimap2, while less effective in these metrics, demonstrate the highest computational efficiency.

## 1 Introduction

Metagenomics is the study of the collective genomes of all microorganisms present in a given microecosystem, offering insights into the composition, diversity, and functions of microbial communities. A more recent technique, High-throughput Chromosome Conformation Capture (Hi-C), enables the study of long-range DNA interactions across the genome in an unbiased, genome-wide manner and can be effectively combined with metagenomics [1]. When integrated with metagenomics, Hi-C enhances traditional shotgun sequencing by providing spatial context to microbial genomes, enabling a deeper understanding of microbial communities. This combined approach not only elucidates the taxonomic composition of a sample but also reveals the interactions of microbial genomes.

The pipeline for generating metagenomic Hi-C (metaHi-C) data has become increasingly more streamlined and standardized. The process begins with collecting environmental samples from the same microbial community, producing shotgun reads via high-throughput sequencing such as Illumina or PacBio and Hi-C reads through paired-end Illumina sequencing. Long-read metaHi-C datasets typically refer to those generated from long-read shotgun libraries using PacBio or Nanopore sequencing, whereas short-read metaHi-C datasets are derived from short-read shotgun libraries using Illumina sequencing. Notably, the difference between long-read metaHi-C datasets and short-read metaHi-C datasets is on the sequencing technology of their shotgun library, not the Hi-C protocol. In Hi-C experiments, DNA is crosslinked to preserve spatial interactions within intact microbial cells, and proximity ligation is performed to create chimeric DNA molecules that link physically adjacent genomic regions [2]. These ligation products are then fragmented, sequenced, and used to infer genomic organization and interactions between different regions. Next, the shotgun reads are assembled into contigs. Hi-C reads are then aligned to these contigs to produce a raw metagenomic Hi-C contact matrix where rows and columns correspond to genomic regions or contigs. Each entry in the matrix represents interaction frequency, reflecting their spatial proximity. This contact matrix can be normalized to correct for biases and is subsequently used for contig binning, a core downstream task in metagenomics where assembled contigs are grouped into metagenome-assembled genomes (MAGs) to recover the structure and diversity of microbial communities de novo. In Hi-C-augmented datasets, the number of read pairs linking two contigs correlates with the probability that they originate from the same genome, making Hi-C contact frequency a strong binning signal [3]. Several pipelines leverage this signal in distinct ways. MetaTOR [4] normalizes raw contacts by the product of contig coverages and clusters the normalized graph with the Louvain algorithm [5] under the Newman–Girvan modularity criterion [6]. HiCBin [7] models experimental bias with HiCzin [8], a normalization method tailored to metaHi-C data, and then clusters with the Leiden algorithm [9]. Bin3C [10] divides counts by the number of restriction sites, applies Knight–Ruiz balancing [11] to obtain a doubly stochastic matrix, and clusters with Infomap [12]. More recently, ImputeCC [13] integrates Hi-C interactions with the discriminative signal from single-copy marker genes to seed and refine bins. In this benchmark, we adopt ImputeCC because it has shown consistent and robust performance on real metagenomic datasets [13].

A critical step in the metagenomic Hi-C analysis pipeline is the alignment of Hi-C reads to assembled contigs. This step directly affects the construction of the raw Hi-C contact matrix. These contacts define contig–contig relationships for genome retrieval and, ultimately, shape the community composition inferred from recovered MAGs [4, 7, 10, 13, 14]. However, aligning Hi-C reads to metagenomic contigs poses unique challenges. First, Hi-C data contains paired reads originating from distant regions, which follow a statistical distribution that differs significantly from that of shotgun libraries. Second, metagenomic datasets include genomes from multiple species, adding complexity to the alignment process. Third, gaps between assembled contigs can lead to lower mapping rates. As a result, general-purpose alignment tools require adjustments to effectively align Hi-C reads to assembled contigs. While there are some benchmark studies evaluating general alignment tools [15, 16], no comprehensive benchmarks exist for metagenomic Hi-C data specifically. Moreover, there is no consensus on a standardized alignment strategy in Hi-C analyses, and various tools have been employed across different studies. For example, Stalder et al [17] used BWA MEM [18], Baudry et al [4] used Bowtie2 [19], Wang et al [20] applied Chromap [21], and Gounot el al [22] leveraged Minimap2 [23]. This variability highlights the need for a systematic benchmark specifically designed to address the unique requirements of alignment tools for metaHi-C data.

## 2 Overview of Alignment Methods for Hi-C data

Alignment methods for metagenomics Hi-C data are essential for processing and analyzing the interactions between metagenomic fragments. These alignment methods aim to map sequencing reads from Hi-C libraries to assembled contigs, facilitating the identification of intra- and inter-contig interactions. In the following, we will briefly review some of the alignment programs that have been used in Hi-C data analysis.

BWA is one of the most widely used alignment tools in genomics due to its efficiency and adaptability to various sequencing data types. It uses the Burrows-Wheeler Transform (BWT) and FM-index to compress the reference genome, enabling fast and memory-efficient alignments [18]. Bowtie2 is another popular aligner for Hi-C data, known for its speed and flexibility. Like BWA, it leverages the BWT and FM-index for efficient alignment yet introduces enhancements for gapped alignments and supports dynamic programming to facilitate local alignment [19]. Chromap and Minimap2 are two other tools employed for Hi-C alignment. Chromap is a recently developed alignment tool [21], designed specifically for chromatin-focused experiments such as Hi-C [24], ChIP-seq [25], and ATAC-seq [26]. It uses a minimizer sketch and split-alignment for efficient and specialized read alignment [21]. Minimap2, originally developed for aligning genomic DNA only, has been extended to map mRNAs as well, making it one of the first RNA-seq aligners specifically designed for long noisy reads [23].

### 2.1 Challenges of Aligning MetaHi-C Reads to Assembled Contigs

Aligning Hi-C reads to metagenomically assembled contigs presents several unique challenges due to the nature of Hi-C data and the complexities of metagenomic assemblies. First, Hi-C data intentionally contains paired reads which can come from very far away genomic regions. Most aligners in paired-end mode operate under the assumption that the paired reads are derived from a single continuous genomic fragment, with an insert size that fits within a known distribution [27]. However, Hi-C ligation products can vary dramatically in size, ranging from 1 bp to hundreds of megabases in terms of linear genomic distance [27]. This makes it challenging to directly apply standard paired-end alignment modes to Hi-C data, as the statistical distribution of paired reads is fundamentally different [27]. Consequently, only a few alignment tools, such as Minimap2, BWA, and Chromap, natively support chimeric alignments and are suitable for Hi-C data.

Secondly, a microbial community often contains hundreds or thousands of microbial organisms, with genome abundance levels varying widely [28]. Additionally, some microbial genomes are highly similar, which can result in ambiguous mapping of Hi-C reads. This similarity, combined with the inherent variability in genome abundance, complicates alignment further, as Hi-C reads can be ambiguously mapped, potentially resulting in low mapping quality of inter-contig relationships.

Lastly, metagenomic assemblies often have gaps between contigs that may not fully represent the complete genomes of all present species, especially in regions with repetitive sequences [29]. When Hi-C reads are aligned to metagenomic assembled contigs, regions near assembly gaps may experience lower alignment accuracy or complete lack of alignment due to missing sequences. These challenges underline the need for alignment tools or robust strategies to effectively map Hi-C reads to metagenomic assemblies.

### 2.2 Selected Alignment Strategies

To evaluate the performance of different alignment methods on metagenomic Hi-C data, we focused seven widely used alignment strategies and tested them on eight environments. The first three strategies—BWA MEM with the -5SP parameter, BWA MEM (P) with default settings, and BWA aln (S) with default settings—are part of the BWA software package, which employs the Burrows-Wheeler Transform (BWT) for string matching. Here, (P) and (S) refer to paired-read and single-read alignment mode, respectively. BWA aln is tailored for short-read alignment. Burton et al [30] used -n 0 on the synthetic yeast metaHi-C dataset, enforcing global alignment with zero mismatches. While effective for the synthetic dataset, we found that the parameter -n 0 is too stringent for real metaHi-C datasets, resulting in notable low mapping rate (Supplementary Table S1). Therefore, in this paper, we opted to use BWA aln with default parameters instead. Compared to BWA aln, BWA MEM is optimized for longer sequences, ranging from 70 base pairs to several megabases. Using the ‘-5SP’ parameter in BWA MEM, pair-end reads are treated as independent single-end reads.

Read pairing and mate-pair rescue functions were deactivated and primary alignments forced to be the alignment with lowest read coordinate (5’ end) [10].

The next two strategies, Bowtie2 (S) with default settings and Bowtie2 (S) with the –very-sensitive-local parameter, originate from the Bowtie2 aligner. Bowtie2 excels at aligning reads ranging from approximately 50 bases to one thousand bases, particularly well-suited for aligning to long genomes [19]. The –very-sensitive-local setting adjusts multiple parameters to maximize sensitivity for local alignment.

The final two strategies, Minimap2 was tested using its default parameters. Minimap2 is a versatile sequence alignment tool capable of aligning DNA or mRNA sequences to large reference databases. Chromap was tested using its Hi-C default parameters. Chromap is specifically designed for ultrafast alignment and preprocessing of high-throughput chromatin profiles. A detailed description of the parameters, settings, and usage examples in Hi-C studies for each strategy is provided in Table 1.

**Table 1.** Summary of the seven alignment strategies evaluated in this study, including their descriptions and usage examples. In the “Methods” column, (P) indicates paired-end alignment mode, while (S) represents single-end alignment mode.

| Methods | Parameter | Description | Usage Example |
| --- | --- | --- | --- |
| BWA MEM (P) | default | BWA maps DNA sequences against a large reference genome. BWA MEM supports long reads and chimeric alignment. | Stalder et al[17], Ivanova et al[31] |
| BWA MEM | -5SP | “-5” forces the alignment to start with 5' end. “SP” skips mate rescue and pairing. | Beyi et al[32], Hwang et al[33], Bickhart et al[34] |
| BWA aln (S) | default | Specifically designed for aligning short DNA sequencing reads (typically up to 100 bp) to a reference genome. | Burton et al[30] |
| Bowtie2 (S) | default | Bowtie 2 is an ultrafast and memory-efficient tool for aligning sequencing reads to long reference sequences. | Kalmart et al[35] |
| Bowtie2 (S) | `-very-sensitive-local` | Performing local alignment and maximizing alignment sensitivity. | Baudry et al[4], Marbouty et al[36] |
| Minimap2 (P) | default | Minimap2 is a versatile sequence alignment program that aligns DNA or mRNA sequences against a large reference database. | Gounot et al[22] |
| Chromp (P) | Hi-C default | Chromap is an ultrafast method for aligning and preprocessing high throughput chromatin profiles. | Wang et al[20] |

## 3 Results

### 3.1 Benchmarking Results for Long-Read MetaHi-C Datasets

We first evaluated the number of inter-contig Hi-C read pairs identified by different alignment strategies on the two long-read MetaHi-C datasets. The corresponding result is shown in Fig. 1. BWA MEM -5SP significantly outperformed other strategies for both long reads datasets (sheep gut and cow rumen), highlighting its superior ability to detect interactions between different genomic regions. In the sheep gut dataset (Fig. 1A), BWA MEM -5SP captures 999,896 inter-contig Hi-C read pairs. BWA MEM default and Minimap2 rank second and third, capturing 677,105 and 653,250 Hi-C read pairs, respectively. In contrast, Chromap had the poorest performance, identifying only 170,415 Hi-C read pairs. In the cow rumen dataset (Fig. 1B), BWA MEM -5SP captures 665,999 inter-contig Hi-C read pairs. BWA MEM default and Chromap follow, detecting 587,641 and 518,855 read pairs. Bowtie2 default demonstrated the lowest performance, identifying only 300,881 Hi-C read pairs. Minimap2 default failed to retain any contigs after minimum MAPQ filtering, resulting in zero usable alignments for downstream analysis.

**Fig. 1.**
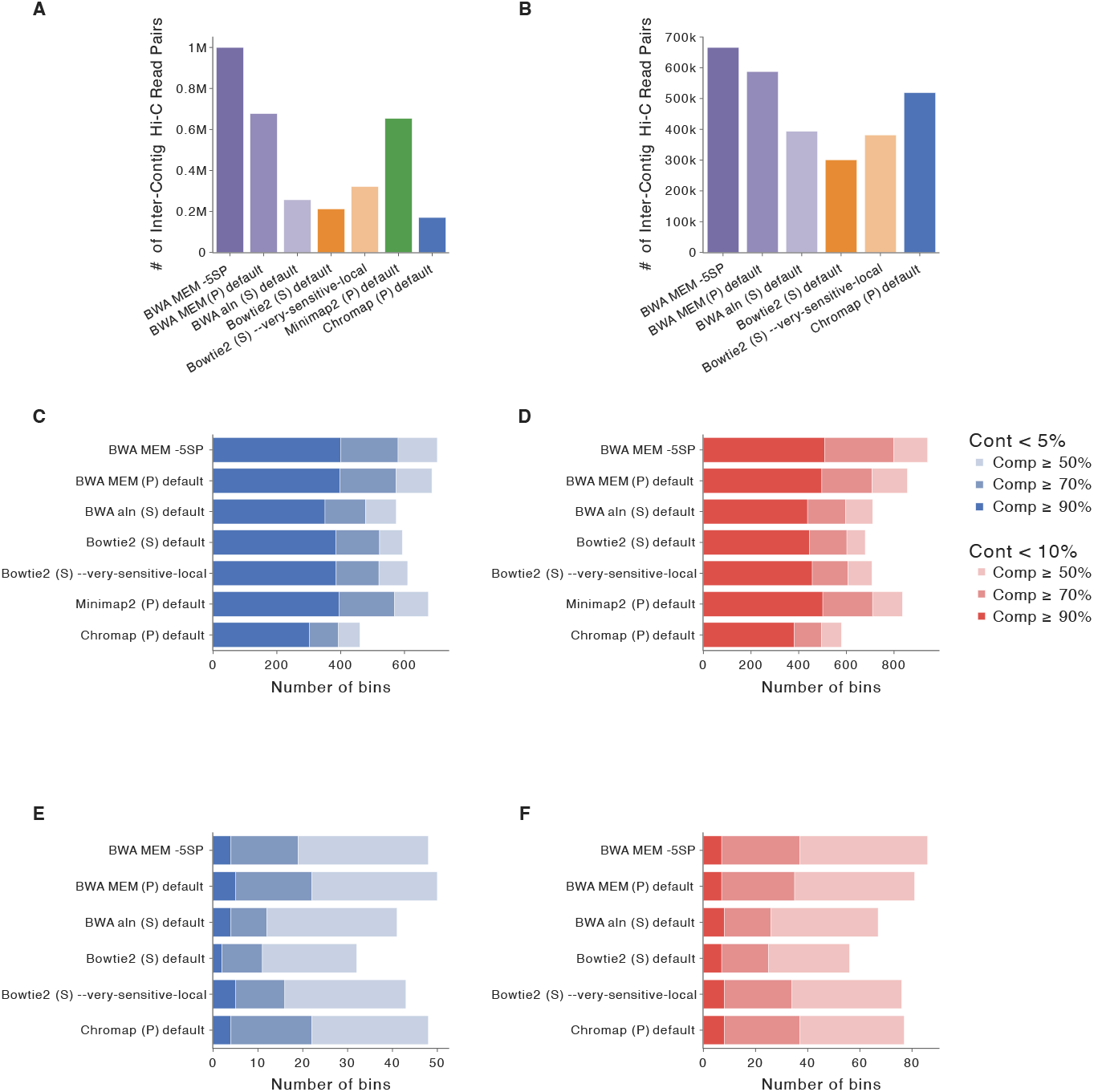
The long-read metagenomics Hi-C datasets benchmarking results. In the legend, “Cont” stands for Contamination, and “Comp” stands for Completeness. **A**: The number of intercontig read pairs captured using different alignment strategies in the sheep gut dataset. **B**: The number of inter-contig read pairs captured using different alignment strategies in the cow rumen dataset. **C**: The number of bins with contamination *<* 5% and completeness ≥50%, 70%, and 90% in the sheep gut dataset. **D**: The number of bins with contamination *<* 10% and completeness ≥50%, 70%, and 90% in the sheep gut dataset. **E**: The number of bins with contamination *<* 5% and completeness ≥ 50%, 70%, and 90% in the cow rumen dataset. **F**: The number of bins with contamination *<* 10% and completeness ≥ 50%, 70%, and 90% in the cow rumen dataset.

From the alignment BAM files, we observed that the mapping rate, defined as the percentage of Hi-C reads successfully aligned to metagenomically assembled contigs, was significantly lower for the cow rumen dataset compared to the sheep gut dataset. For instance, using BWA MEM -5SP, the cow rumen dataset achieved a mapping rate of 51.55%, compared to 85.53% for the sheep gut dataset. This discrepancy arises from differences in the shotgun libraries: the cow rumen dataset was derived from error-prone PacBio long reads, whereas the sheep gut dataset was based on highly accurate HiFi reads. As a result, mapping accurate Hi-C reads to an error-prone long-read assembly leads to a lower mapping rate.

As for the binning results based on CheckM2, BWA MEM -5SP achieved the highest total number of bins with contamination below 5% and completeness ≥50%. Moreover, in the sheep gut dataset (Fig. 1C), it identified the most bins meeting the stringent threshold of ≥90% completeness and contamination below 5%. BWA MEM default also showed good performance similar to BWA MEM -5SP, ranked second. In contrast, Chromap demonstrated the weakest performance, yielding the lowest binning results across all metrics. A similar trend was observed when the contamination is below 10% (Fig. 1D). In the cow rumen dataset (Fig. 1E), BWA MEM -5SP and Chromap default produced the second-highest number of bins with contamination below 5% and completeness ≥50%, while BWA MEM default demonstrated the best performance. Under more relaxed criteria (Fig. 1F), BWA MEM -5SP outperformed all aligners, generating the most bins with contamination below 10% and completeness ≥50%, followed by BWA MEM default and Chromap default.

To further investigate how different aligners influence the inferred taxonomic profiles of the microbial community, we performed bin-level taxonomic annotation of bins with ≥50% completeness and *<* 10% contamination using GTDB-Tk [37] (see Subsection 5.4). We first summarized, for each alignment strategy, the number of unique taxa (family, genus, species) recovered from GTDB-Tk annotations of the resulting MAG sets. In the cow rumen dataset (Supplementary Tables S10), BWA-MEM default recovered the highest number of unique species (97) followed by BWA MEM -5SP (92), while Minimap2 recovered the fewest (62). In the sheep gut dataset (Supplementary Tables S10), both BWA MEM -5SP and BWA MEM default performed best, each identifying 261 unique species, whereas Chromap recovered the fewest (194). To illustrate the overlap and divergence in taxonomic profiles, we selected one high-performing aligner (BWA-MEM -5SP), one moderately performing aligner (Chromap default), and one low-performing aligner (Bowtie2 –very-sensitive-local) to generate Venn diagrams at both the genus and species levels (Supplementary Fig. 21). In the cow rumen dataset, BWA MEM -5SP identified 7 unique species not recovered by the other two aligners, whereas in the sheep gut dataset it recovered 40 unique genera that the other two did not identify. These results highlight that aligner choice can substantially impact final biological resolution in the long-read datasets.

In conclusion, BWA MEM -5SP consistently outperforms other alignment strategies in detecting inter-contig Hi-C read pairs and achieving superior binning quality for long-read metaHi-C datasets. In contrast, the performance of other strategies varies depending on the environmental context.

### 3.2 Benchmarking Results for Short-Read MetaHi-C Datasets

#### 3.2.1 Synthetic Yeast Dataset

We compared the number of inter-contig Hi-C read pairs identified by each alignment strategies shown in Fig. 2A. Among the tested strategies, the BWA-based approaches outperformed Bowtie2, Minimap2, and Chromap in detecting inter-contig read pairs. Notably, BWA MEM with the -5SP parameter identified the highest number of intercontig Hi-C read pairs. BWA MEM with default parameters and BWA aln ranked second and third, respectively.

**Fig. 2.**
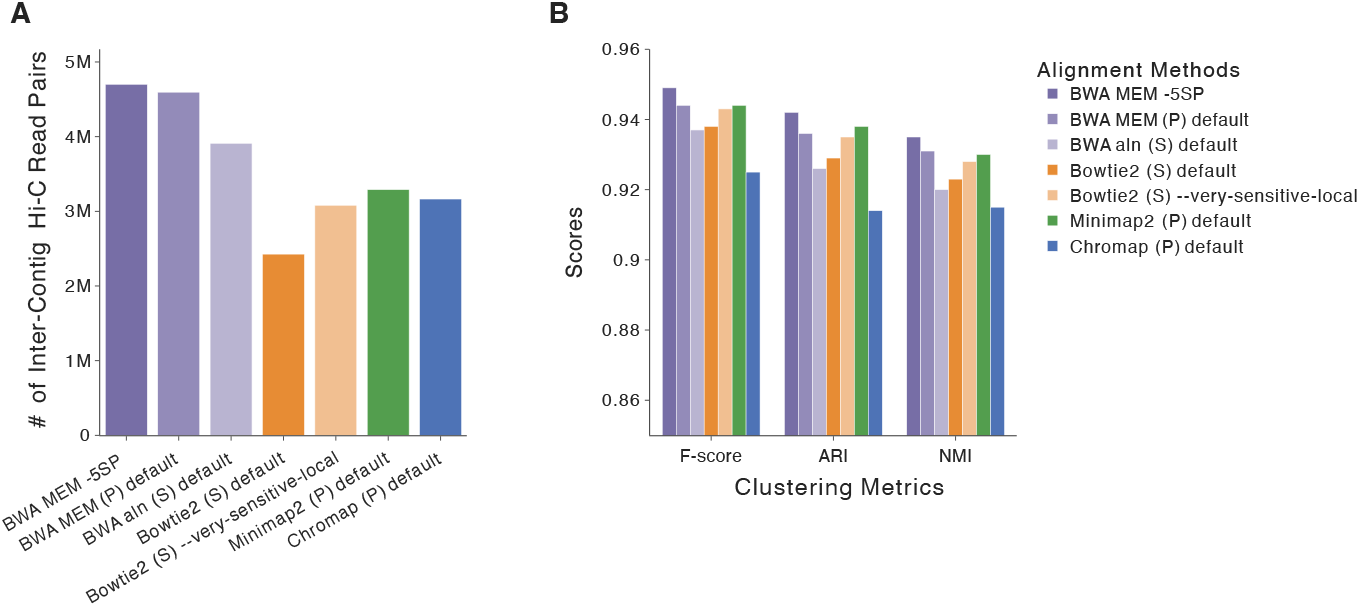
The synthetic yeast dataset benchmarking results. **A**: The number of inter-contig read pairs captured and **B**: the clustering performance metrics (F-score, ARI, and NMI) for different alignment strategies.

Leveraging the known species identities of all contigs in this synthetic dataset, we further evaluated clustering performance using the Fowlkes-Mallows score (F-score), Adjusted Rand Index (ARI), and Normalized Mutual Information (NMI) as shown in Fig. 2B. While all seven alignment strategies exhibited robust performance, BWA MEM -5SP achieved the highest scores across all three metrics, followed by Minimap2 and BWA MEM with default settings.These results indicate that the clusters produced by BWA MEM -5SP align most accurately with the true species-level clustering of contigs. Overall, based on the number of inter-contig read pairs identified and clustering performance, BWA MEM -5SP demonstrates the best performance among all the alignment strategies tested in this synthetic yeast dataset.

#### 3.2.2 Real Short-Read MetaHi-C Datasets

Using evaluation metrics similar to those applied to long-read metaHi-C datasets, we first compare the number of inter-contig Hi-C read pairs captured by different alignment strategies. For single-dataset environments, such as bovine and wastewater, BWA MEM -5SP consistently detects the highest number of inter-contig read pairs. Similarly, for multi-dataset environments, including hydrothermal mats, pigs, and human gut environments, BWA MEM -5SP achieves the best performance in most datasets, demonstrating its effectiveness across a diverse range of metagenomic Hi-C datasets.

For the bovine dataset in Fig. 3A, BWA MEM -5SP captured the highest number of inter-contig Hi-C read pairs, totaling 3,514,606. BWA MEM default ranked second with 2,700,382 inter-contig read pairs. In contrast, Bowtie2 and Chromap performed similarly, capturing approximately one-fifth of the read pairs detected by BWA MEM -5SP. For the wastewater dataset in Fig. 3B, BWA MEM -5SP identified 20,529,313 inter-contig Hi-C read pairs. BWA MEM default again ranked second with 13,032,526 read pairs, followed by Minimap2, which detected 11,282,105 read pairs. In contrast, strategies such as Bowtie2 (default and very-sensitive-local modes) detected fewer than 5,000,000 inter-contig read pairs, yielding the lowest count among all strategies. For the human gut environment in Fig. 3C, BWA-MEM -5SP demonstrated consistent superiority, detecting the highest number of inter-contig Hi-C read pairs in 23 out of 24 datasets. BWA MEM default ranked second, identifying a substantial number of inter-contig Hi-C read pairs across most datasets, though consistently performing below BWA MEM -5SP. Its performance showed a strong correlation with that of BWA MEM -5SP; when the performance of BWA MEM -5SP declined, BWA MEM default also exhibited a significant drop. Among the other strategies, Minimap2 exhibited moderate performance, while Chromap consistently detected the lowest number of inter-contig Hi-C read pairs. A similar trend was observed in the hydrothermal mats and pigs environments (Supplementary Figures 1 and 2), where BWA-MEM -5SP consistently detected the highest number of inter-contig Hi-C read pairs, followed by BWA-MEM with default settings. This pattern is expected because paired-end insert-size modeling is ill-suited to Hi-C. In default mode, BWA-MEM estimates an insert-size distribution and rescues mates that conform to it, which can suppress true long-range ligation products. Running BWA-MEM with -5SP disables mate rescue and insert-size-based pairing, effectively aligning mates independently. Across all datasets, this configuration yields more inter-contig Hi-C pairs than the default mode (Fig. 3), consistent with the long-range nature of Hi-C contacts.

**Fig. 3.**
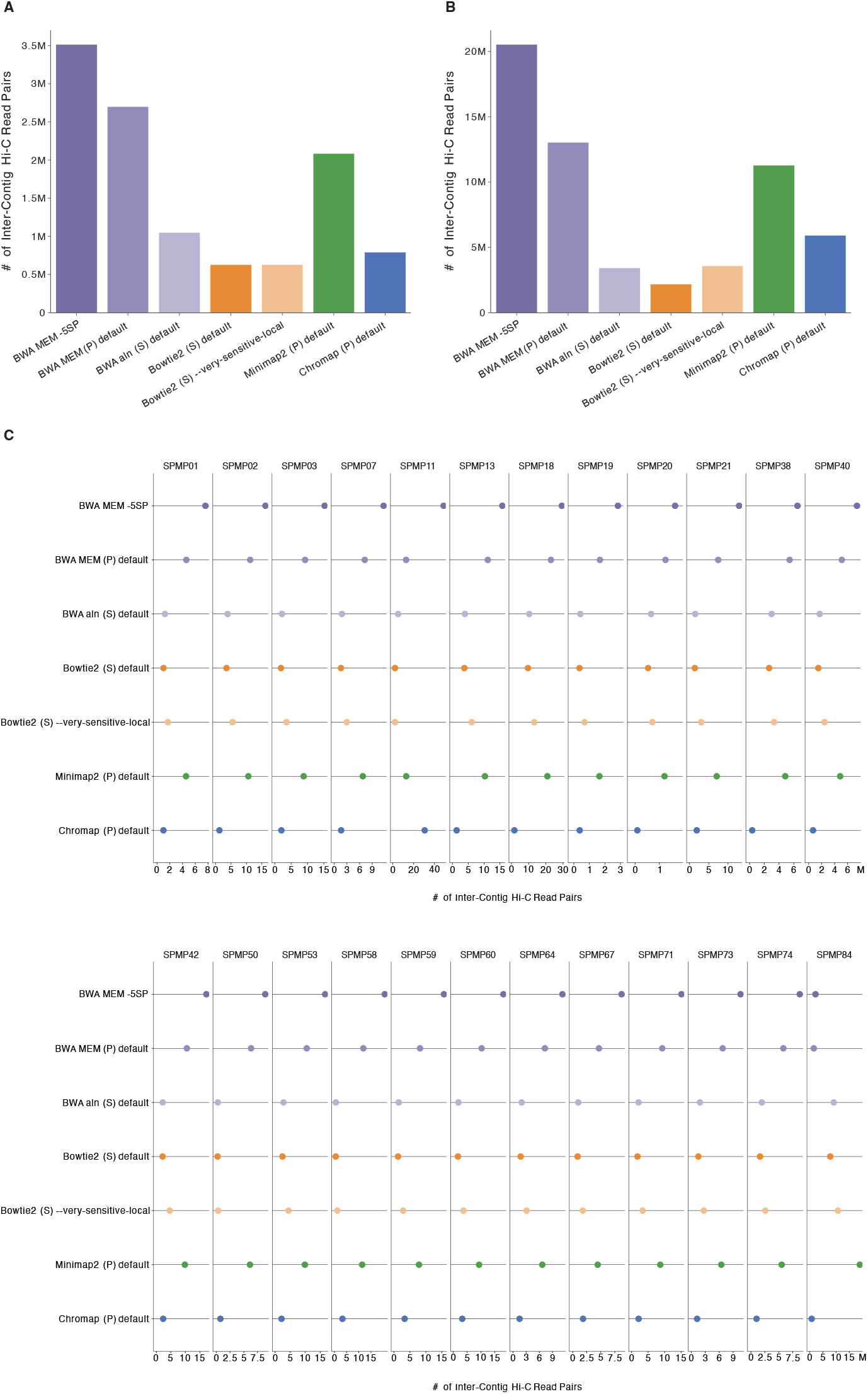
Comparison of inter-contig read pairs captured by different alignment strategies across various datasets. **A**: Bovine environment. **B**: Wastewater environment. **C**: Human gut environment.

BWA MEM -5SP excels not only in identifying inter-contig Hi-C read pairs but also in producing the best binning results among all alignment strategies. For the bovine dataset (Fig. 4A), BWA MEM -5SP achieved the highest total number of bins with contamination below 5% and completeness ≥50%, followed by BWA MEM default and Minimap2 default. Conversely, Chromap identified the fewest bins meeting these criteria. A similar trend was observed when the contamination threshold was set to below 10% (Fig. 4B). For the wastewater dataset (Fig. 4C), BWA-MEM -5SP generated nearly twice as many bins with contamination *<* 5% and completeness ≥50% compared to BWA aln. It ranked first overall, generating the most bins with contamination *<* 5% and completeness ≥ 90%. Minimap2 and BWA-MEM default ranked second and third. The trend is similar when contamination *<* 10% (Fig. 4D). For the hydrothermal mats environment (Fig. 5A), BWA MEM -5SP achieved the best binning results in 6 out of 10 datasets, generating the highest number of bins with contamination *<* 5% and completeness ≥90%. Even in the three datasets where it was not the top-performing aligner, BWA MEM -5SP still ranked highly, outperforming most other strategies. BWA MEM -5SP demonstrated the best performance in bins with contamination *<* 10% and completeness ≥90% across 8 out of 10 datasets when using a more lenient contamination threshold (Fig. 5B). In the pigs environment (Fig. 5C), BWA MEM -5SP maintained its top performance, producing the highest number of bins with contamination *<* 5% and completeness ≥90% in 16 out of 20 datasets. Notably, it generated over 65 bins meeting these criteria for dataset CV2 1 and delivered strong results in other datasets such as CV7 2 and CV10 1. BWA MEM default also performed well, ranking second in overall binning quality. While it produced substantial numbers of bins with contamination *<* 5% and completeness ≥90% across most datasets, it lagged behind BWA-MEM -5SP. When contamination is less than 10%, a silimar trend was observed in (Fig. 5D), followed by BWA MEM default and Minimap2 default. For the human gut environment (Fig. 5E), BWA MEM -5SP consistently outperformed other strategies in 11 datasets, producing the highest number of bins with contamination *<* 5% and completeness ≥90%. BWA MEM default closely trails BWA MEM -5SP in several datasets such as SPMP03, SPMP19 and SPMP40. Minimap2 also showed strong performance; it performs equally or better than BWA MEM -5SP in 15 datasets. Using the contamination *<* 10% and completeness ≥90% threshold (Fig. 5F), BWA MEM -5SP, Minimap2 default, and BWA MEM default are the three best-performing aligners.

**Fig. 4.**
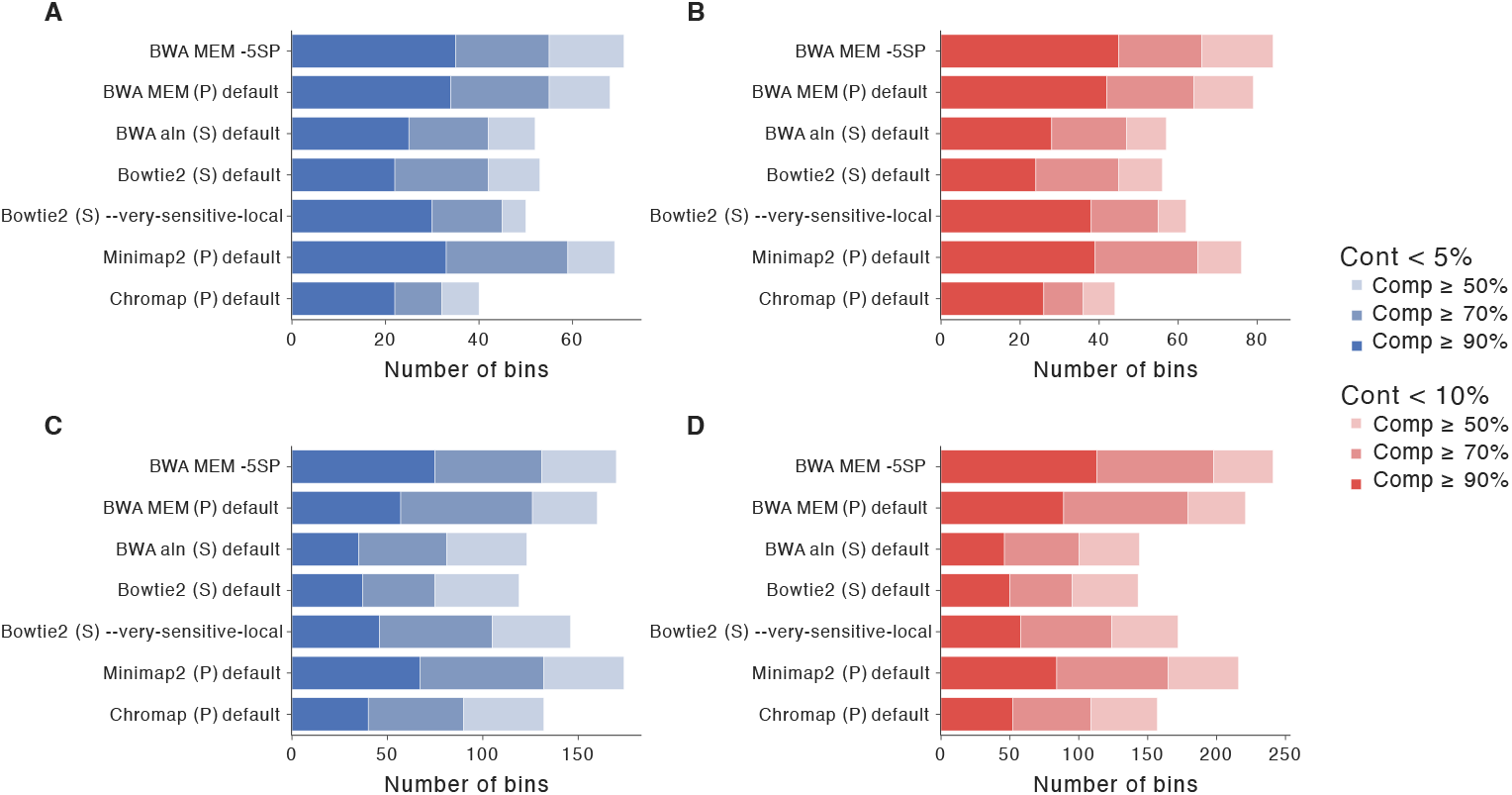
Comparison of binning results between bovine and wastewater datasets. In the legend, “Cont” stands for Contamination, and “Comp” stands for Completeness. **A**: The number of bins with contamination *<* 5% and completeness ≥50%, 70%, and 90% in the bovine dataset. **B**: The number of bins with contamination *<* 10% and completeness ≥50%, 70%, and 90% in the bovine dataset. **C**: The number of bins with contamination *<* 5% and completeness ≥90% in the wastewater dataset. **D**: The number of bins with contamination *<* 10% and completeness ≥90% in the wastewater dataset.

**Fig. 5.**
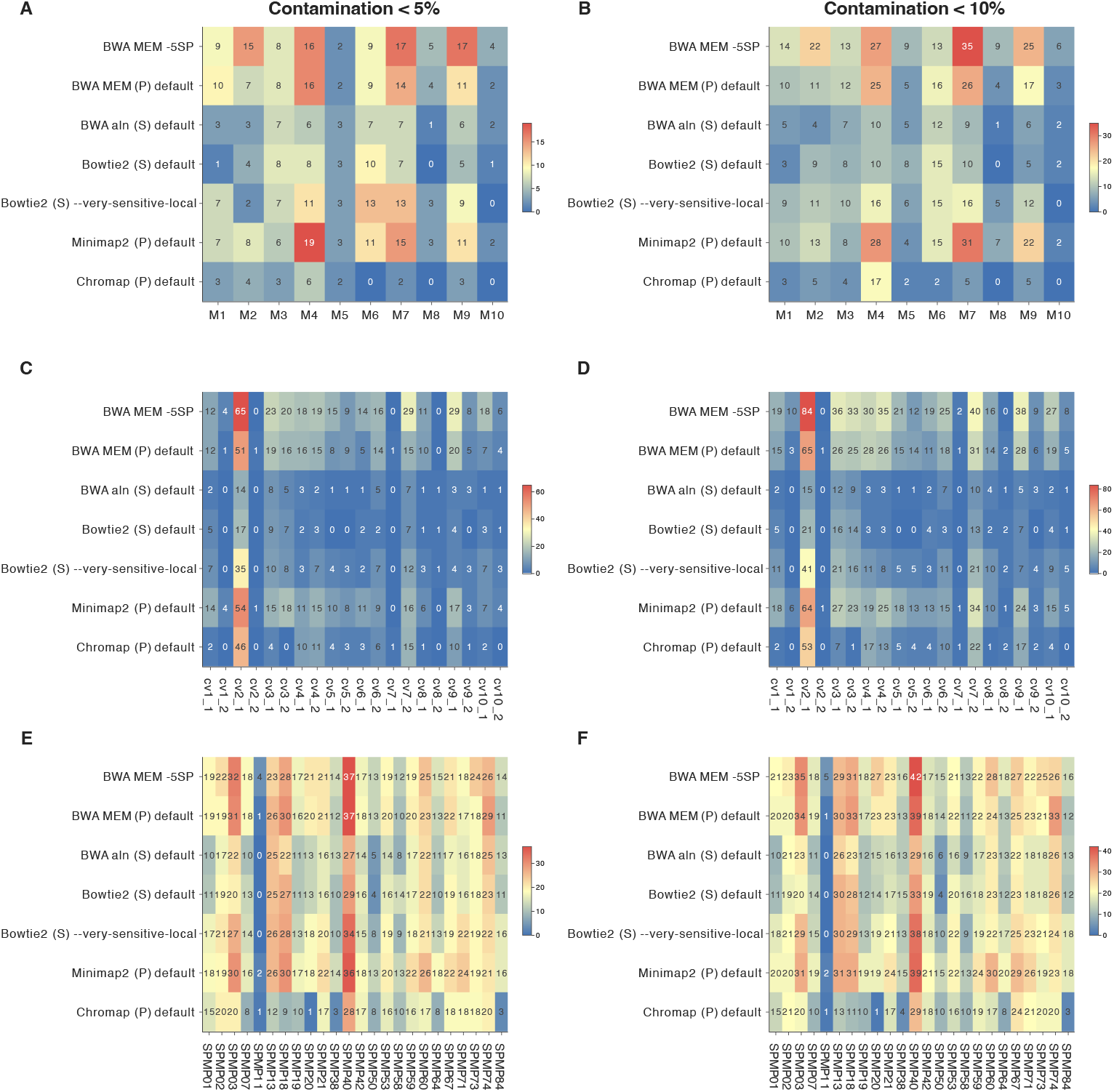
Comparison of binning results across different alignment strategies and datasets. **A**: The number of bins with contamination *<* 5% and completeness ≥90% in the hydrothermal mats environment. **B**: The number of bins with contamination *<* 10% and completeness ≥90% in the hydrothermal mats environment. **C**: The number of bins with contamination *<* 5% and completeness ≥ 90% in the pigs environment. **D**: The number of bins with contamination *<* 10% and completeness ≥ 90% in the pigs environment. **E**: The number of bins with contamination *<* 5% and completeness ≥ 90% in the human gut environment. **F**: The number of bins with contamination *<* 10% and completeness ≥ 90% in the human gut environment.

We then performed the same annotation analysis as in the long-read metaHi-C datasets on the short-read metaHi-C environments. For the hydrothermal mats, pigs, and human gut environments, we selected one representative dataset per environment. Specifically, we chose the dataset with the smallest combined number of bases from the metagenomic and Hi-C libraries. We summarized the number of unique taxa (family, genus, species) represented in the MAG sets produced under each alignment strategy (Supplementary Tables S5-S9). For instance, in the bovine dataset, MAGs derived with BWA MEM -5SP spanned 44 unique genera, whereas those from Chromap with the Hi-C default only spanned 19. We visualized shared and unique taxa across strategies with Venn diagrams (Supplementary Fig. 22). The overall pattern is consistent with the long-read datasets, indicating that aligner choice can substantially influence the diversity and number of taxa recovered in downstream analyses for short-read metaHi-C datasets as well.

### 3.3 Computational Time Complexity

All the datasets were ran on a High-Performance-Cluster (HPC) with a xeon-2640356 16-core CPU node and 100 GB of memory. Fig. 6 presents the execution time of the alignment strategies across different datasets. For the long-read sheep gut dataset (Fig. 6A), Bowtie2 –very-sensitive-local had the highest execution time, while Minimap2 and Chromap exhibited the lowest runtime. Other strategies, including BWA MEM variations, demonstrated intermediate runtimes. In the cow rumen dataset (Fig. 6B), Minimap2 was the fastest followed by Chromap default and Bowtie2 default strategies. In short-read yeast dataset (Fig. 6C), Minimap2 had the highest computational efficiency followed by BWA MEM strategies. For the bovine short-read dataset (Fig. 6D), Bowtie2 –very-sensitive-local again displayed the highest runtime, while Minimap2 achieved the fastest runtime. For the wastewater dataset (Fig. 6E), Minimap2 was the most efficient followed by Chromap and BWA MEM strategies. For the hydrothermal mats environment with 10 datasets (Fig. 6F), Bowtie2 –verysensitive-local exhibited greater variability and higher median runtimes compared to Minimap2 and Chromap. In the pigs environment with 20 datasets (Fig. 6G), Minimap2 demonstrated the shortest runtime while Bowtie2 –very-sensitive-local showed longer runtime. Lastly, in the human gut environment with 24 datasets (Fig. 6H), Minimap2 again achieved the most efficient runtime followed by BWA MEM strategies. Overall, Minimap2 and Chromap are the most computationally efficient strategies, though their performance exhibits some variability across different environments. BWA MEM strategies exhibited intermediate to relatively low runtimes, while Bowtie2 –very-sensitive-local tended to be the slowest.

**Fig. 6.**
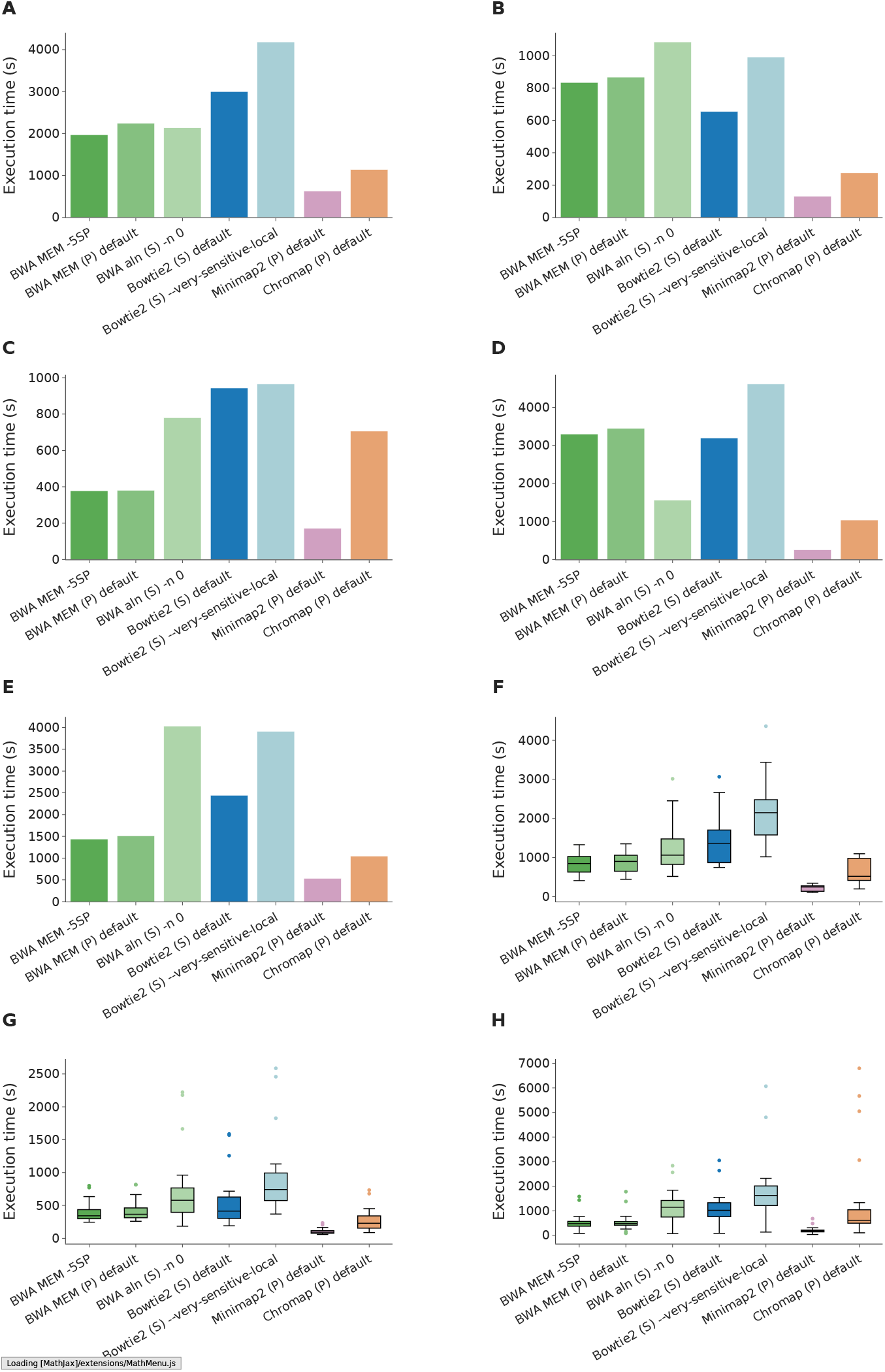
Runtime comparison of alignment strategies across various datasets. **A**: Sheep gut environment. **B**: Cow rumen environment. **C**: Synthetic yeast environment **D**: Bovine environment. **E**: Wastewater environment. **F**: Hydrothermal mats environment. **G**: Pigs environment. **H**: Human gut environment.

## 4 Discussion

In this work, we focused on evaluating seven commonly used alignment strategies in metagenomics Hi-C: BWA MEM -5SP, BWA MEM (P) default, BWA aln (S) default, Bowtie2 (S) default, Bowtie2 (S) –very-sensitive-local, Minimap2 (P) default, and Chromap (P) Hi-C default. These alignment strategies are often used to map Hi-C reads to assembled metagenomic contigs. This is an important step in metagenomic Hi-C analysis as it directly affects the quality of the raw metagenomic Hi-C contact matrix. However, there is currently no consensus on the optimal alignment strategy to use for this task. To address this gap, we benchmarked these alignment strategies using one synthetic dataset and seven real datasets. Our results show that while the performance of the alignment strategies varies, BWA MEM -5SP consistently outperforms all others across different environments. BWA MEM default closely follows, demonstrating strong but slightly lower performance compared to BWA MEM -5SP. The superior performance of BWA MEM -5SP is mainly due to the -SP flag, which disables read pairing and mate-pair rescue functions and treats paired reads as independent single-end reads. In default paired-end mode, the aligner tries to estimate an insert size and preferentially rescue pairs that align close to each other, which can suppress the detection of true long-range Hi-C contacts. In addition, the -5 flag takes the lowest coordinate (5’ end) as the primary alignment. This simplifies downstream data processing for metaHi-C pipelines, since they typically rely on the primary alignment [4, 7, 10, 38–40]. In comparison, Minimap2 and Chromap, though efficient, are more likely to miss or misplace short chimeric Hi-C hits, leading to lower sensitivity in this context. In comparison, Minimap2 and Chromap, though efficient, are more likely to miss or misplace short chimeric Hi-C hits, leading to lower sensitivity in this context. Although Chromap and Minimap2 perform less effectively in the evaluation metrics, they demonstrate the highest computational efficiency across most environments. Similarly, BWA MEM strategies also exhibit relatively high computational efficiency.

In addition to benchmarking the seven core alignment strategies, which use default or widely used settings from prior Hi-C studies, we also investigated the impact of alternative parameter settings and varying MAPQ thresholds (Supplementary Figures 14 to 20). Specifically, we evaluated variants including Bowtie2 (S) –very-sensitive, Minimap2 (P) with the -x sr preset, and Chromap with -e 8 under the default Hi-C preset. Overall, the recommended configurations of these aligners generally outperformed their alternative parameterizations. As for the mapping-quality cutoff, in addition to the widely used default cutoff of 30 [14, 40–42], we evaluated a threshold of 20 reported in prior work [43] and a more lenient cutoff of 10 (Supplementary Figures 14 to 20). As the MAPQ threshold increases, the number of inter-contig Hi-C read pairs decreases. Across MAPQ thresholds, BWA MEM -5SP remains the best-performing aligner in most cases. When varying MAPQ for BWA MEM -5SP and evaluating bins with completeness ≥90% and contamination *<* 5%, MAPQ 30 yields the most bins in five of seven datasets, whereas the bovine and pig datasets peak at MAPQ 10. This indicates that MAPQ selection can influence binning performance. Therefore, in practice, although MAPQ 30 is a good and widely used choice, testing a small set of mapping-quality cutoffs (e.g., 10, 20, 30) can be considered when computational resources permit, with the final value chosen to optimize binning performance according to CheckM2 or other selected criteria.

Finally, several reference properties also modulate alignment performance. Contig fragmentation (shorter contigs with more termini) reduces effective unique k-mer context and increases multi-mapping; local aligners also more often partially align reads near contig ends. Both effects lower MAPQ [19, 44]. Strain-level redundancy likewise creates competing targets and reduces MAPQ; in a sensitivity analysis, dereplicating contigs with CD-HIT-EST [45] modestly increased inter-contig contacts in both a short-read (wastewater) and a long-read (cow rumen) metaHi-C datasets, but produced no substantial difference in downstream binning performance (Supplementary Note 2; Supplementary Tables S3 and S4). In the future, since different assemblers may produce contigs with different qualities, such as differences in contiguity and base accuracy, it is worth exploring how different assemblers influence metaHi-C analyses. Moreover, our benchmarks are tailored to metagenomic Hi-C, where multi species complexity, fragmented assemblies, and ambiguous mapping dominate. It remains to be established whether our observations, for example aligning mates independently to avoid insert size assumptions, generalize to human focused Hi-C alignment, where analyses are performed against a nearly complete, high quality reference genome. Therefore, a systematic human focused Hi-C alignment benchmark is an important direction for future work.

## 5 Materials and Methods

### 5.1 Datasets

To evaluate the alignment performance of different strategies, we chose one synthetic yeast dataset and several real metagenomic Hi-C datasets derived from diverse environments. Among the real datasets, five environments are based on short-read shotgun libraries, while the sheep gut and cow rumen environments originate from long-read shotgun libraries. All of them have short-read Hi-C libraries. Notably, for every dataset, Hi-C reads were generated with Illumina sequencing. We label a dataset ‘long-read metaHi-C’ when its shotgun library was generated using third-generation sequencing and ‘short-read metaHi-C’ when its shotgun library was generated using second-generation sequencing. An overview of these datasets is provided in Table 2, and the data is available in Additional file 2.

**Table 2.** Descriptions of eight environments used in the experiments. In the “Long Read / Short Read” column, “Long Read” denotes long-read metaHi-C datasets, and “Short Read” represents short-read metaHi-C datasets. The “Restriction Enzymes” column specifies the restriction enzymes used for constructing each corresponding Hi-C library.

| Environment | Description | Long read / Short read | Restriction enzymes |
| --- | --- | --- | --- |
| Sheep gut | A fecal sample was taken from a young (<1 year old) wether lamb of the Katahdin breed. | Long | Sau3AI and MluCI |
| Cow Rumen | Rumen contents from one multiparous Holstein cow housed at the University of Wisconsin, Madison, campus were sampled via rumen cannula. | Long | Sau3AI and MluCI |
| Synthetic yeast | Mixture of yeasts, consisting of 13 yeast species. | Short | NcoI and HindIII |
| Bovine | Large pooled samples collected from cows with lesions of DD at slaughterhouses. | Short | Sau3AI and MluCI |
| Wastewater | The wastewater sample was collected in October 2017 at the Moscow WWTP in Idaho (USA). | Short | Sau3AI and MluCI |
| Hydrothermal mats | Microbial mat samples were collected during a research expedition. | Short | Sau3AI and MluCI |
| Pigs | Pig fecal samples came from five conventional pig farms and five organic pig farms. | Short | HpyCH4IV |
| Human gut | Fecal samples were collected from individuals aged 48 to 76 as part of the Singapore Platinum Metagenomes Project (SPMP). | Short | Sau3AI and MluCI |

For the sheep gut long-read metaHi-C dataset [34], it includes PacBio circular consensus sequencing (CCS), or HiFi, long-read libraries and illumina short-read Hi-C libraries. The dataset resulted in 255 Gbp from CCS long-read libraries and 107 million read pairs from Hi-C libraries. The cow rumen long-read metaHi-C dataset [46] includes PacBio uncorrected long-read libraries and Hi-C libraries. The error-prone PacBio long reads were generated using the PacBio RSII and PacBio Sequel platforms, totaling 52 Gbp. The Hi-C libraries were sequenced on an Illumina HiSeq 2000 at 80 bp, producing 63.4 million Hi-C reads.

The synthetic yeast dataset [30] comprises a mixture of 13 yeast species (Supplementary Table S2). The raw shotgun libraries consisted of 85.7 million read pairs at 101 bp, while the raw Hi-C libraries contained 81 million read pairs at 100 bp. The bovine dataset [32] provided a total of 21 Gb of sequence data for the shotgun library and 16.8 Gb of Hi-C data, both sequenced using the Illumina HiSeq platform. For the wastewater dataset [17], both Hi-C and shotgun metagenomic libraries were pooled and sequenced using two lanes of the Illumina HiSeq 4000 platform. The shotgun metagenomic library yielded 269,312,499 read pairs, while the Hi-C library produced 95,284,717 read pairs. While the environments discussed above each contain a single dataset, the next three environments include multiple datasets within each. In the hydrothermal mats environment [33], which includes 10 different metaHi-C datasets, both Hi-C and shotgun libraries in each dataset were sequenced on a single lane of an Illumina NovaSeq S4 system, yielding a total of 1.8 billion shotgun read pairs and 1.5 billion Hi-C read pairs, respectively. The sizes of the Hi-C libraries range from 29.1 Gbp to 70 Gbp. For the pigs environment [35], which comprises 20 datasets, both Hi-C and shotgun libraries were sequenced on the Illumina HiSeq 4000 platform. The shotgun libraries generated a total of 3.5 billion reads, while the Hi-C libraries produced 3.1 billion reads with sizes ranging from 7.1 Gbp to 26.9 Gbp. Lastly, the human gut environment [22], consisting of 24 datasets, was generated using paired-end sequencing on the Illumina HiSeq 4000 platform for both metagenomic and Hi-C libraries. The metagenomic library produced a total of 256 Gbp, while the Hi-C library yielded 564 Gbp. Hi-C library sizes range from 2.4 Gbp to 87 Gbp, whereas shotgun library sizes range from 6 Gbp to 23 Gbp. The restriction enzymes used for constructing Hi-C libraries in each environment are listed in Table 2, while a summary of dataset sizes is provided in Additional file 3.

### 5.2 Data Preprocessing and Assembly

We processed all datasets using a standardized cleaning procedure with BBTools [47]. First, we used BBDuk (v39.06) from BBTools to trim adapters and perform quality filtering through k-mers on paired-end sequencing reads for both Hi-C and WGS data with parameters ‘ktrim=r k=23 mink=11 hdist=1 minlen=50 tpe tbo’. Then, we applied additional quality trimming to sequencing reads with parameters ‘trimq=10 qtrim=r ftm=5 minlen=50’. To eliminate potential biases and remove low-quality bases from library preparation, we further trimmed the first 10 bases from the 5’ end of each Hi-C and WGS sequencing read (ftl=10). Lastly, we utilized Clumpify (v39.06) from BBTools to deduplicate paired-end Hi-C reads with default parameters. For the short reads datasets, the WGS reads were assembled into contigs using Megahit [48] (v1.2.9) with parameters ‘–min-contig-len 1000 –k-min 21 –k-max 141 –k-step 12 –merge-level 20,0.95’. For the two long reads datasets, we used the contigs provided in the original studies. Specifically, for the sheep gut dataset, Bickhart et al [34] assembled the Pacbio long reads using metaFlye (v2.7 ‘—meta —pacbio-hifi’) and deposited at https://doi.org/10.5281/zenodo.5228989 under the file `‘flye.v29.sheep_gut.hifi.250g.fasta.gz’`. For the cow rumen dataset, PacBio raw reads were assembled by Bickhart et al [46] using Canu (v1.6+101 changes (r8513) with parameters ‘-trim-assemble genomeSize=5m oeaMemory=32 redMemory=32 correctedErrorRate=0.035’), followed by two rounds of polishing with Illumina data via Pilon. The final assembly is available at https://figshare.com/articles/usda_pacbio_second_pilon_indelsonly_fa_gz/8323154.

### 5.3 Alignment

For each dataset, we align paired-end Hi-C reads to the assembled contigs from the same sample using seven strategies. With Bowtie2 (v2.3.5.1), we employed two approaches: (1) single-read mode with default parameters and (2) single-read mode with the –very-sensitive-local option to enable local alignment using the very sensitive preset. For BWA (v0.7.17-r1188), we used three strategies: (1) the BWA aln algorithm in single-read mode with default parameters, (2) the BWA MEM algorithm in paired-read mode with default parameters, and (3) the BWA MEM algorithm in paired-read mode with the -5SP parameter. In addition to those, we applied two more tools: Minimap2 (v2.26-r1175) in paired-read mode with default parameters and Chromap (v0.2.6-r490) with default settings. Following each alignment strategy, the output alignment files were converted to sorted BAM files using SAMtools [49] (v1.18).

### 5.4 Downstream Analysis

After alignment, we removed unmapped reads with Samtools [50] and supplied the contig assembly (FASTA) and the alignments (BAM) to NormCC, the normalization module of the MetaCC pipeline [40] (v1.2.0). By default, NormCC filters Hi-C alignments using a minimum MAPQ threshold of 30. The retained alignments are used to construct the raw Hi-C contact matrix, which is then normalized to eliminate systematic biases and produce the final normalized contacts. NormCC requires the restriction enzyme(s) to be specified via the -e option, which we set according to each dataset: for the synthetic yeast dataset, -e NcoI -e HindIII; for the pig datasets, -e HpyCH4IV; and for the other real datasets, -e MluCI -e Sau3AI, matching the enzymes used during library preparation. All other parameters were left at their defaults. Leveraging the normalized Hi-C contact matrix generated from NormCC, we further used ImputeCC [13] (v1.0.0) with default parameters to bin the contigs for all real Hi-C datasets. ImputeCC is not applicable to synthetic yeast dataset because it relies on single-copy marker genes of prokaryotes [13]. Instead, we clustered contigs from the normalized Hi-C contact matrix using the Leiden graph clustering algorithm [9] with default parameters.

We selected one representative dataset from each environment for taxonomic annotation of the bins with CheckM2 completeness ≥50% and contamination *<* 10%. For environments with only one dataset, we use that dataset directly. For environments containing multiple datasets, we chose the one with the smallest combined number of bases from the metagenomic and Hi-C libraries. Taxonomic annotation was performed using GTDB-Tk [37] (v2.5.2) with default parameters (database release 226). We then calculated the number of unique families, genera, and species identified for each aligner. The detailed results are provided in Additional File 6.

### 5.5 Evaluation Criteria

To begin, we assess the performance of alignment strategies by comparing their ability to capture inter-contig contacts. Inter-contig contacts are interactions between sequences located on different contigs. Since metagenomics Hi-C experiments are designed to capture contig-to-contig relationships, the number of inter-contig contacts serves as a practical proxy for alignment sensitivity. Notably, we do not adjudicate the biological accuracy of individual inter-contig contacts, which cannot be established without ground truth on the real datasets. Instead, this metric should be regarded as reflecting the strength of the Hi-C signal linking contigs, consistent with prior metaHi-C work [33, 51, 52].

Besides counting the number of inter-contig contacts, we further investigate the impact of alignment on downstream analyses, such as binning. We use CheckM2 [53] (v1.0.2) to report bin completeness with contamination thresholds below 5% and 10%. CheckM2 employs a machine learning-based approach that leverages a larger set of features to estimate the completeness of genome bins, providing a detailed assessment of how much of the genome has been recovered. Additionally, it uses a gradientboosting model to evaluate the level of contamination in genome bins, suggesting that the genome bin may contain DNA from more than one organism [53].

For the synthetic yeast dataset, unlike the real Hi-C datasets, the ground truth for both intra-species Hi-C contacts and spurious inter-species Hi-C contacts was known (Supplementary Note 1). Since ImputeCC relies on single-copy marker genes of prokaryotes [13], we cannot use ImputeCC to optimize the contig binning process for the yeast dataset. Instead, we directly applied the Leiden graph clustering algorithm with default parameters to the normalized Hi-C contact matrix, clustering contigs into draft genomic bins. To evaluate the performance of contig clustering across different alignment strategies, we computed Fowlkes-Mallows score (F-score), Adjusted Rand Index (ARI), and Normalized Mutual Information (NMI). These metrics provided a comprehensive evaluation of clustering accuracy and alignment performance.

The Fowlkes-Mallows score (F-score) is a metric used to evaluate the similarity between two clusterings, often applied in clustering validation tasks. It is defined as the geometric mean of precision and recall. The score is calculated as follows:

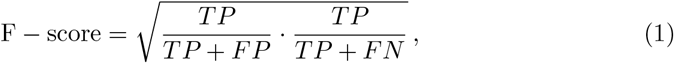

where *TP* is true positive, *FP* is false positive, and *FN* is false negative. The F-score ranges from 0 to 1, where a score closer to 1 indicates higher similarity between the two clusterings, reflecting better clustering performance.

The Rand Index (RI) is a metric used to measure the similarity between two clusterings by considering the consistency of sample pair assignments. It evaluates how well the predicted clustering agrees with the true clustering by comparing pairwise relationships across all samples.

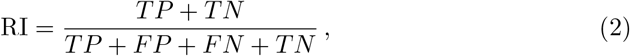

where *TN* is true negative.

The Adjusted Rand Index is a correction of the Rand Index. While the Rand Index evaluates the similarity between two clusterings, it can be biased by the number of clusters or the size of the dataset, leading to inflated scores even for random clusterings. The ARI corrects this by incorporating the Expected Rand Index, which accounts for the similarity expected by chance. It is defined as follows:

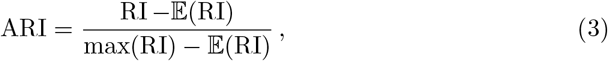

Where 𝔼 (RI) is the Expected Rand Index.

The Normalized Mutual Information is another metric to assess the similarity between two different cluster assignments, providing a score that indicates how much information is shared between the two clusterings. A value closer to 1 means the two clusters are more similar. Let U and V denote the sets of true class labels and predicted cluster labels, respectively. Define the entropy of a label set S as

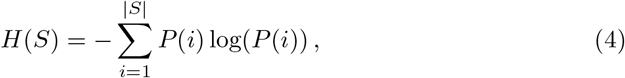

where 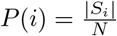 is the probability of an object in class *S*_*i*_.

The Mutual Information (MI) between U and V is calculated by:

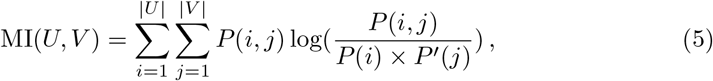

where 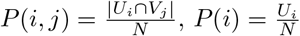, and 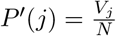.

Therefore, the Normalized Mutual Information is

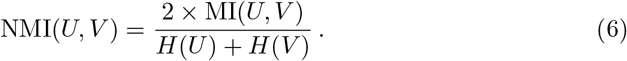

## Declarations

## Acknowledgements

The research is partially funded by NSF grant EF-2125142. Y.D. is partially supported by the University of Texas Systems STAR Program. We thank Dr. Heng Li (author of BWA and Minimap2) and Dr. Li Song (author of Chromap) for their valuable insights and discussions on the BWA, Minimap2, and Chromap aligners.

## Author contributions

Y.D. and F.S. conceived the study and designed the research. Y.W. and Y.D. performed the computational analyses and drafted the manuscript. W.Z. and J.H. collected and analyzed the human gut datasets. All authors contributed to revising and finalizing the manuscript.

## Consent for publication

All authors have approved the manuscript for submission.

## Data Availability

The data used in this study can be accessed and downloaded from Additional file 2.

## Code Availability

The benchmarking code is publicly available at https://github.com/yuw053/MetaHi-C_Benchmark.git.

## Ethics approval and consent to participate

Not applicable.

## Supplementary information

### Additional file 1

Supplementary notes, figures and tables (PDF 377 KB). Identifying the species of assembled contigs on the synthetic yeast dataset, supplementary table S1-S13 and supplementary figures 1-22 supporting the manuscript.

### Additional file 2

Overview of Raw Data Across Environments (xlsx 19 KB). Sheet 1 includes data for the pigs environment, while Sheet 2 contains data for the human gut environment.

Sheet 3 provides data for the bovine, hydrothermal mats, wastewater, sheep gut, cow rumen, and synthetic yeast environments.

### Additional file 3

The size of shotgun and Hi-C libraries from raw metaHi-C datasets (xlsx 11 KB).

### Additional file 4

Overview of runtime Across datasets (xlsx 14 KB). This table provides the detailed runtime for each alignment strategy applied to each dataset.

