## Supplemental File 1 for "Benchmarking Alignment Strategies for Hi-C Reads in Metagenomic Hi-C Data"

### Supplementary Notes

##### Supplementary Note 1: Identifying the species of assembled contigs on the synthetic yeast dataset

The reference genomes for all 16 yeast strains representing 13 yeast species in the synthetic yeast sample were downloaded from Supplementary Table 2. Since the analysis was conducted at the species level, the genomes of four strains belonging to the same species (Saccharomyces cerevisiae—FY, CEN.PK, RM11-1A, and SK1) were combined into a single reference genome. Subsequently, all contigs were aligned to the reference genomes of the 13 known species using BLAST [1] with the parameters '-perc_identity 95 -evalue 1e-30 -word_size 50'. The true species identity of each assembled contig was inferred based on its alignment to the reference genome of a specific species.

##### Supplementary Note 2: The effect of dereplicating metagenomic WGS assembled contigs

To further evaluate the effect of strain-level redundancy, we followed the strategy used in Hwang et al. [2] and applied CD-HIT-EST [3] dereplication (-c 0.95 -aS 0.85) to the assembled contigs in order to remove highly similar sequences. We tested this on one short-read dataset (wastewater) and one long-read dataset (cow rumen) using the BWA MEM –5SP aligner. The results are presented in Supplementary Tables 3 and 4. In the waste water dataset, the number of inter-contig read pairs have been increased from 20,503,126 to 21,174,130. A similar trend has also been observed in the cow rumen dataset. This behavior is expected, as removing redundant contigs reduces ambiguity in alignment and improves the recovery of valid Hi-C contacts across distinct genomic regions. However, we observed no substantial difference in downstream binning performance.

### Supplementary Tables

**Table 1 Comparison of Mapping Rates Among BWA Alignment Strategies.** This table presents the mapping rates of three BWA alignment strategies across three datasets. Notably, BWA aln -n 0 exhibits a substantially lower mapping rate compared to the other strategies.

| Dataset | BWA MEM -5SP | BWA MEM default | BWA aln -n 0 |
|---|---|---|---|
| Sheep gut | 85.53 | 91.01 | 54.135 |
| Bovine | 57.78 | 63.5 | 14.05 |
| Pig cv7_2 | 79.85 | 87.21 | 20.995 |

**Table 2 The species list in the synthetic yeast dataset.**

| Genus | Species | Strain in sample | Reference strain |
|---|---|---|---|
| *Saccharomyces* | *cerevisiae* | FY4H | FY |
| *Saccharomyces* | *cerevisiae* | CEN.PK | CEN.PK |
| *Saccharomyces* | *cerevisiae* | RM11-1A | RM11-1A |
| *Saccharomyces* | *cerevisiae* | SK1 | SK1 |
| *Saccharomyces* | *paradoxus* | YDG613 | YDG613 |
| *Saccharomyces* | *mikatae* | FM356 | IFO 1815 |
| *Saccharomyces* | *kudriavzevii* | FM527 | IFO 1802 |
| *Saccharomyces* | *bayanus var. uvarum* | YZB5-113 | CBS 7001 |
| *Naumovozyma* | *castellii* | 4310 | NRRL Y-12630 |
| *Lachancea* | *waltii* | Kwaltii ura3 | NRRL Y-8285 |
| *Lachancea* | *kluyveri* | FM628 | CBS 3082 |
| *Kluyveromyces* | *lactis* | MW98-8C | NRRL Y-1140 |
| *Kluyveromyces* | *wickerhamii* | Y-8286 | UCD 54-210 |
| *Ashbya* | *gossypii* | WT | ATCC 10895 |
| *Scheffersomyces* | *stipitis* | Y-11545 | CBS 6054 |
| *Pichia* | *pastoris* | JC308 | GS115 |

**Table 3 Impact of Contig Dereplication on Mapping and Binning in the Wastewater Dataset.** This table compares the number of inter-contig Hi-C read pairs and binning quality before and after dereplicating assembled contigs using CD-HIT-EST.

| **Metric** | **Before** | **After** |
|---|---|---|
| # of inter-contig read pairs | 20,529,313 | 21,174,130 |
| # bins with contamination < 5% | 172 | 170 |
| # bins with contamination < 10% | 243 | 236 |

**Table 4 Impact of Contig Dereplication on Mapping and Binning in the Cow Rumen Dataset.** This table compares the number of inter-contig Hi-C read pairs and binning quality before and after dereplicating assembled contigs using CD-HIT-EST.

| **Metric** | **Before** | **After** |
|---|---|---|
| # of inter-contig read pairs | 665,999 | 682,788 |
| # bins with contamination < 5% | 48 | 44 |
| # bins with contamination < 10% | 86 | 82 |

**Table 5 Taxonomic Annotation Results in the Human Gut Dataset (SPMP11).** The number of unique families, genera, and species recovered by each alignment strategy.

| Alignment Strategy | # of unique family | # of unique genus | # of unique species |
|---|---|---|---|
| Bowtie2 (S) default | 1 | 1 | 1 |
| Bowtie2 (S) –very-sensitive-local | 1 | 2 | 2 |
| Bowtie2 (S) –very-sensitive | 1 | 1 | 1 |
| BWA MEM -5SP | 7 | 15 | 16 |
| BWA MEM (P) default | 6 | 9 | 9 |
| Chromap (P) default -e 8 | 6 | 11 | 11 |
| Chromap (P) default | 6 | 13 | 13 |
| Minimap2 (P) default | 6 | 10 | 9 |
| Minimap2 (P) -x sr | 5 | 7 | 5 |

**Table 6 Taxonomic Annotation Results in the Hydrothermal Mats Dataset (M8).** The number of unique families, genera, and species recovered by each alignment strategy.

| Alignment Strategy | # of unique family | # of unique genus | # of unique species |
|---|---|---|---|
| Bowtie2 (S) default | 11 | 13 | 9 |
| Bowtie2 (S) –very-sensitive-local | 13 | 16 | 13 |
| Bowtie2 (S) –very-sensitive | 10 | 11 | 6 |
| BWA aln (S) default | 11 | 12 | 8 |
| BWA MEM -5SP | 20 | 26 | 21 |
| BWA MEM (P) default | 17 | 22 | 15 |
| Chromap (P) default -e 8 | 10 | 11 | 6 |
| Chromap (P) default | 11 | 12 | 7 |
| Minimap2 (P) default | 17 | 22 | 14 |
| Minimap2 (P) -x sr | 16 | 22 | 18 |

**Table 7 Taxonomic Annotation Results in the Pig Gut Dataset (cv7_1).** The number of unique families, genera, and species recovered by each alignment strategy.

| Alignment Strategy | # of unique family | # of unique genus | # of unique species |
|---|---|---|---|
| Bowtie2 (S) default | 4 | 5 | 4 |
| Bowtie2 (S) –very-sensitive-local | 7 | 7 | 11 |
| Bowtie2 (S) –very-sensitive | 3 | 4 | 4 |
| BWA aln (S) default | 2 | 2 | 1 |
| BWA MEM -5SP | 12 | 19 | 29 |
| BWA MEM (P) default | 11 | 13 | 19 |
| Chromap (P) default -e 8 | 3 | 3 | 1 |
| Chromap (P) default | 5 | 5 | 4 |
| Minimap2 (P) default | 10 | 12 | 17 |
| Minimap2 (P) -x sr | 7 | 9 | 12 |

**Table 8 Taxonomic Annotation Results in the Bovine Dataset.** The number of unique families, genera, and species recovered by each alignment strategy.

| Alignment Strategy | # of unique family | # of unique genus | # of unique species |
|---|---|---|---|
| Bowtie2 (S) default | 21 | 29 | 18 |
| Bowtie2 (S) –very-sensitive-local | 25 | 34 | 21 |
| Bowtie2 (S) –very-sensitive | 22 | 30 | 18 |
| BWA aln (S) default | 22 | 30 | 18 |
| BWA MEM -5SP | 30 | 44 | 28 |
| BWA MEM (P) default | 28 | 39 | 25 |
| Chromap (P) default -e 8 | 17 | 20 | 15 |
| Chromap (P) default | 17 | 19 | 15 |
| Minimap2 (P) default | 26 | 40 | 23 |
| Minimap2 (P) -x sr | 27 | 40 | 24 |

**Table 9 Taxonomic Annotation Results in the Wastewater Dataset.** The number of unique families, genera, and species recovered by each alignment strategy.

| Alignment Strategy | # of unique family | # of unique genus | # of unique species |
|---|---|---|---|
| Bowtie2 (S) default | 61 | 109 | 124 |
| Bowtie2 (S) –very-sensitive-local | 69 | 126 | 145 |
| Bowtie2 (S) –very-sensitive | 60 | 106 | 129 |
| BWA aln (S) default | 60 | 105 | 124 |
| BWA MEM -5SP | 80 | 165 | 194 |
| BWA MEM (P) default | 80 | 161 | 185 |
| Chromap (P) default -e 8 | 63 | 116 | 125 |
| Chromap (P) default | 66 | 126 | 134 |
| Minimap2 (P) default | 76 | 146 | 169 |
| Minimap2 (P) -x sr | 76 | 150 | 175 |

**Table 10 Taxonomic Annotation Results in the Cow Rumen Dataset.** The number of unique families, genera, and species recovered by each alignment strategy.

| Alignment Strategy | # of unique family | # of unique genus | # of unique species |
|---|---|---|---|
| Bowtie2 (S) default | 16 | 38 | 67 |
| Bowtie2 (S) –very-sensitive-local | 17 | 45 | 86 |
| BWA aln (S) default | 16 | 42 | 76 |
| BWA MEM -5SP | 19 | 50 | 92 |
| BWA MEM (P) default | 19 | 50 | 97 |
| Chromap (P) default | 19 | 45 | 83 |
| Minimap2 (P) -x sr | 15 | 35 | 62 |

**Table 11 Taxonomic Annotation Results in the Sheep Dataset.** The number of unique families, genera, and species recovered by each alignment strategy.

| Alignment Strategy | # of unique family | # of unique genus | # of unique species |
|---|---|---|---|
| Bowtie2 (S) default | 86 | 295 | 228 |
| Bowtie2 (S) –very-sensitive-local | 88 | 315 | 233 |
| Bowtie2 (S) –very-sensitive | 86 | 295 | 226 |
| BWA aln (S) default | 87 | 297 | 234 |
| BWA MEM -5SP | 96 | 357 | 261 |
| BWA MEM (P) default | 96 | 349 | 261 |
| Chromap (P) default -e 8 | 82 | 269 | 194 |
| Chromap (P) default | 78 | 265 | 194 |
| Minimap2 (P) default | 94 | 335 | 257 |
| Minimap2 (P) -x sr | 95 | 336 | 254 |

### Supplementary Figures

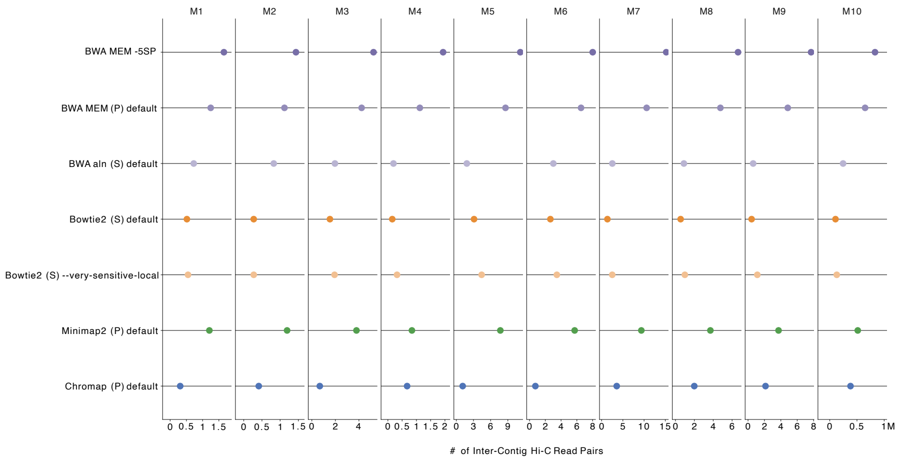


**Fig. 1 Comparison of inter-contig read pairs captured by different alignment methods in hydrothermal mats dataset.** Results for datasets M1 to M10.

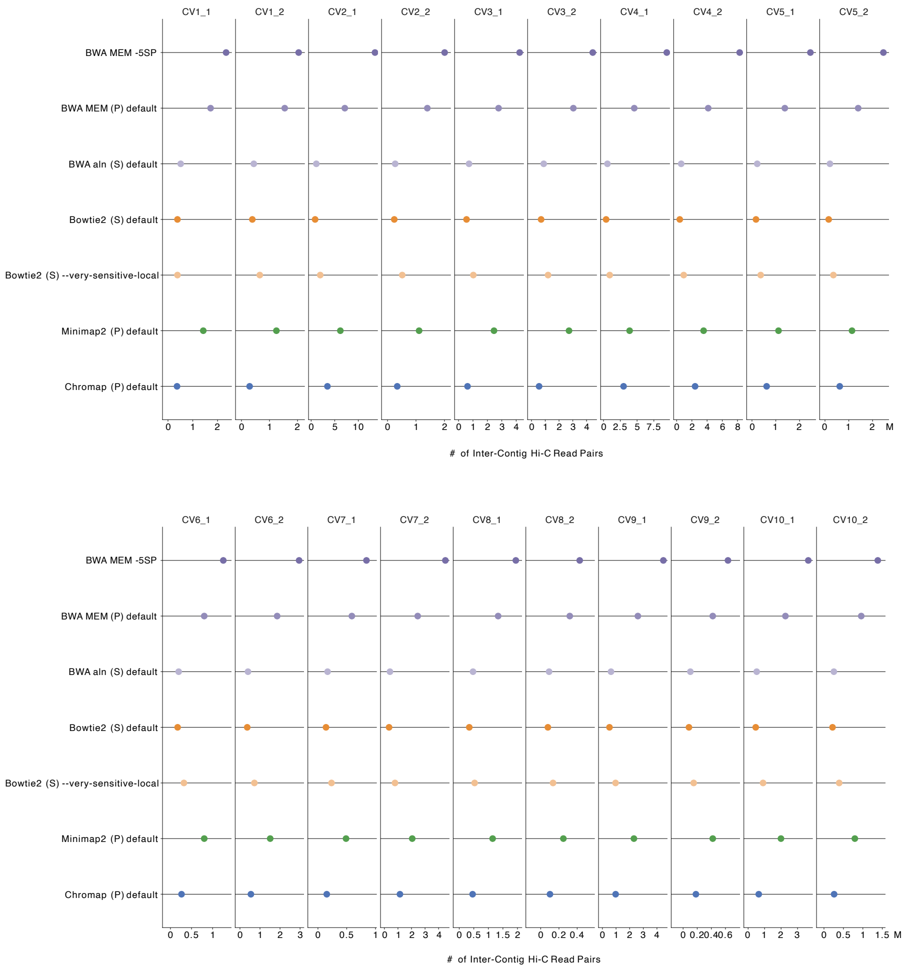


**Fig. 2 Comparison of inter-contig read pairs captured by different alignment methods in pigs dataset.** Results for datasets CV1_1 to CV10_2.

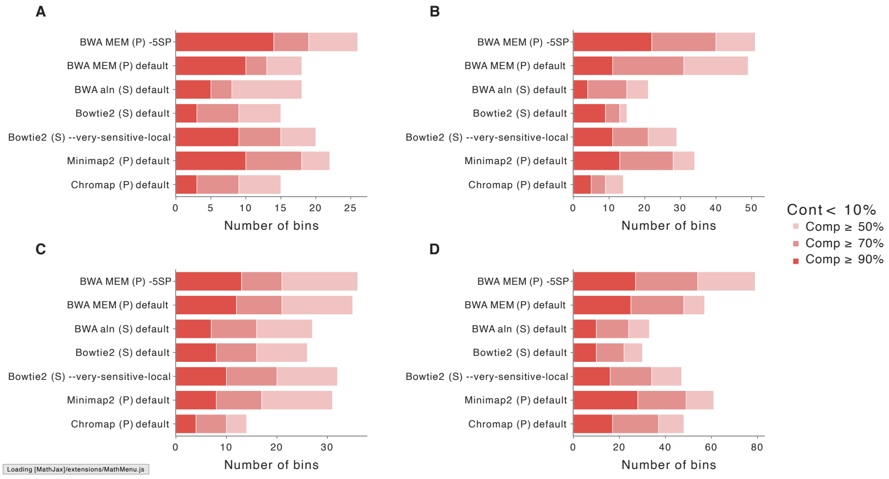


**Fig. 3 Binning results for hydrothermal mats datasets.** The legend abbreviations are: "Cont" for Contamination and "Comp" for Completeness. Panels **A-D** show the number of bins with contamination $< 10\%$ and completeness thresholds of $\geq 50\%$, 70%, and 90% for the hydrothermal mats datasets M1 (**A**), M2 (**B**), M3 (**C**), and M4 (**D**).

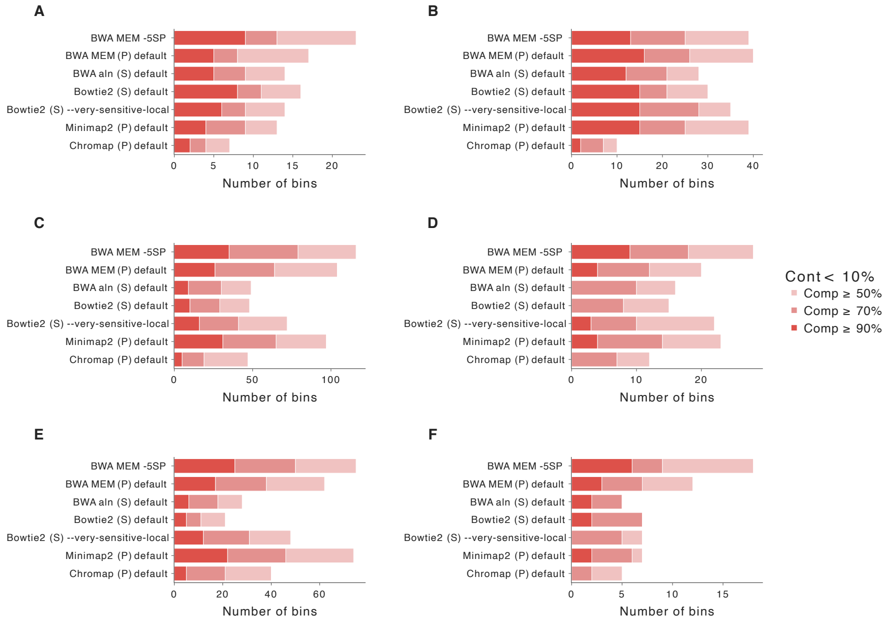


**Fig. 4 Binning results for hydrothermal mats datasets.** The legend abbreviations are: "Cont" for Contamination and "Comp" for Completeness. Panels **A-F** show the number of bins with contamination $< 10\%$ and completeness thresholds of $\geq 50\%$, 70%, and 90% for the hydrothermal mats datasets M5 (**A**), M6 (**B**), M7 (**C**), M8 (**D**), M9 (**E**), and M10 (**F**).

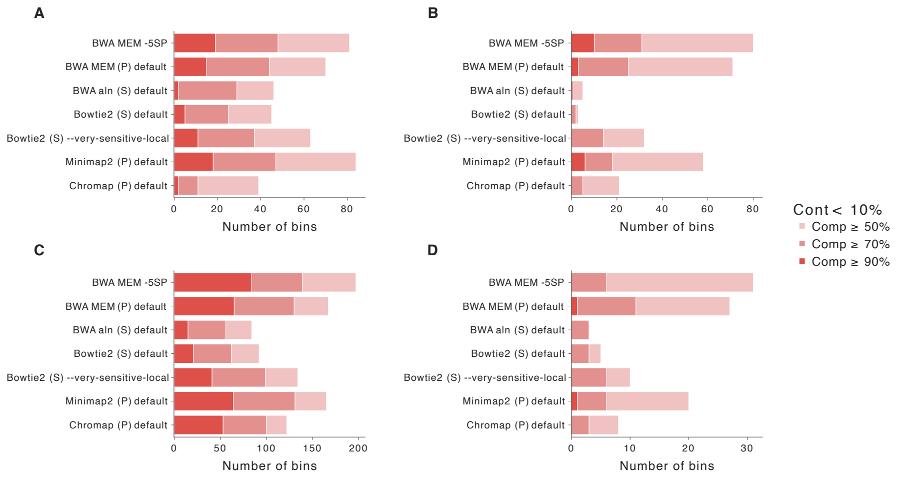


**Fig. 5 Binning results for pigs datasets.** The legend abbreviations are: "Cont" for Contamination and "Comp" for Completeness. Panels **A-D** show the number of bins with contamination $<$ 10% and completeness thresholds of $\geq$ 50%, 70%, and 90% for the pig datasets cv1_1 (**A**), cv1_2 (**B**), cv2_1 (**C**), and cv2_2 (**D**).

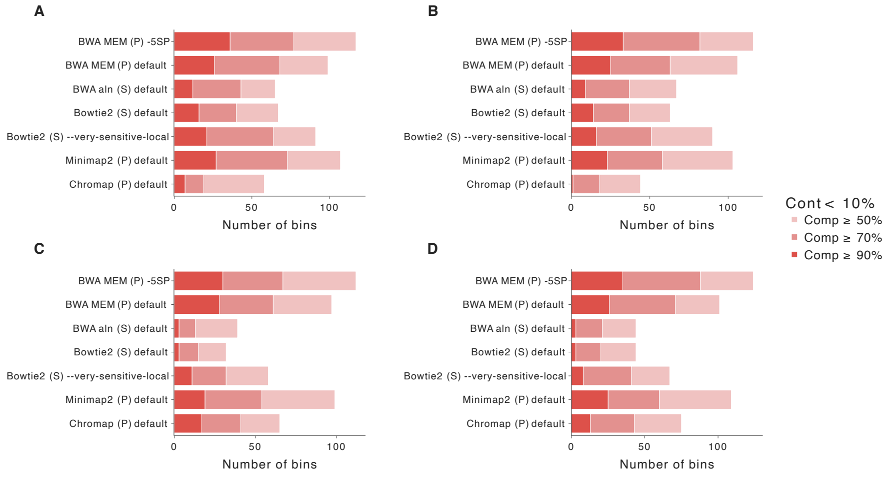


**Fig. 6 Binning results for pigs datasets.** The legend abbreviations are: "Cont" for Contamination and "Comp" for Completeness. Panels **A-D** show the number of bins with contamination $<$ 10% and completeness thresholds of $\geq$ 50%, 70%, and 90% for the pig datasets cv3_1 (**A**), cv3_2 (**B**), cv4_1 (**C**), and cv4_2 (**D**).

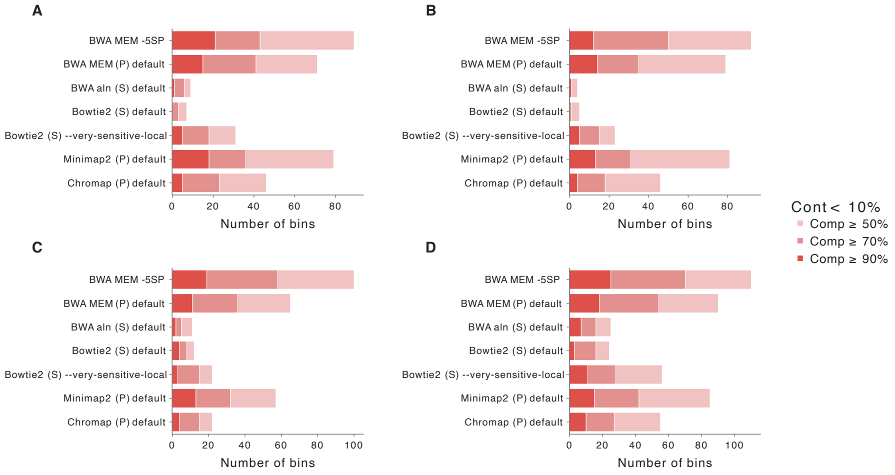


**Fig. 7 Binning results for pigs datasets.** The legend abbreviations are: "Cont" for Contamination and "Comp" for Completeness. Panels **A-D** show the number of bins with contamination $<$ 10% and completeness thresholds of $\geq$ 50%, 70%, and 90% for the pig datasets cv5_1 (**A**), cv5_2 (**B**), cv6_1 (**C**), and cv6_2 (**D**).

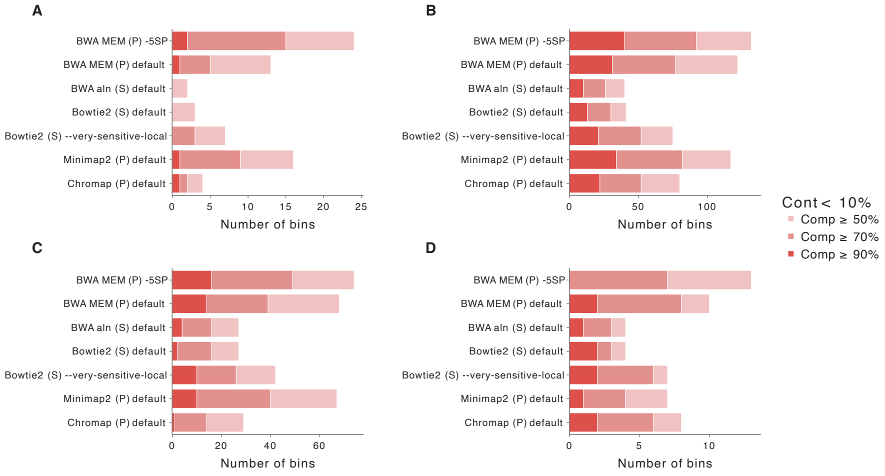


**Fig. 8 Binning results for pigs datasets.** The legend abbreviations are: "Cont" for Contamination and "Comp" for Completeness. Panels **A-D** show the number of bins with contamination $<$ 10% and completeness thresholds of $\geq$ 50%, 70%, and 90% for the pig datasets cv7_1 (**A**), cv7_2 (**B**), cv8_1 (**C**), and cv8_2 (**D**).

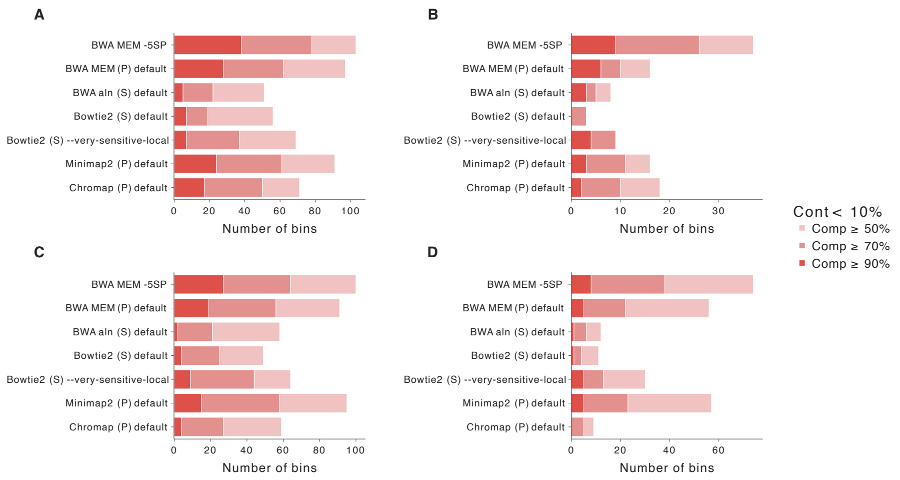


**Fig. 9 Binning results for pigs datasets.** The legend abbreviations are: "Cont" for Contamination and "Comp" for Completeness. Panels **A-D** show the number of bins with contamination < 10% and completeness thresholds of ≥ 50%, 70%, and 90% for the pig datasets cv9_1 (**A**), cv9_2 (**B**), cv10_1 (**C**), and cv10_2 (**D**).

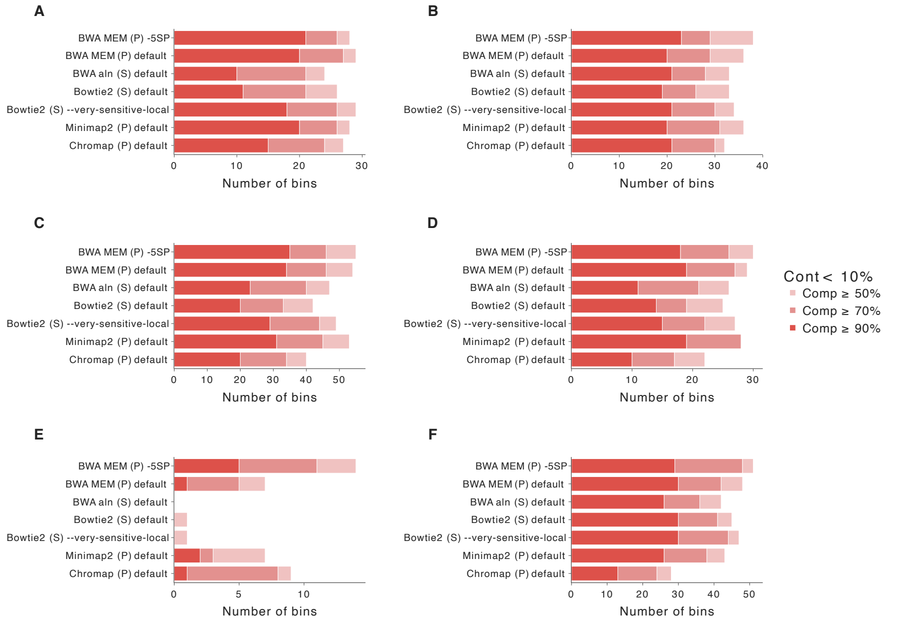


**Fig. 10 Binning results for human gut datasets.** The legend abbreviations are: "Cont" for Contamination and "Comp" for Completeness. Panels **A-F** show the number of bins with contamination < 10% and completeness thresholds of ≥ 50%, 70%, and 90% for the human gut datasets SPMP01 (**A**), SPMP02 (**B**), SPMP03 (**C**), SPMP07 (**D**), SPMP11 (**E**), and SPMP13 (**F**).

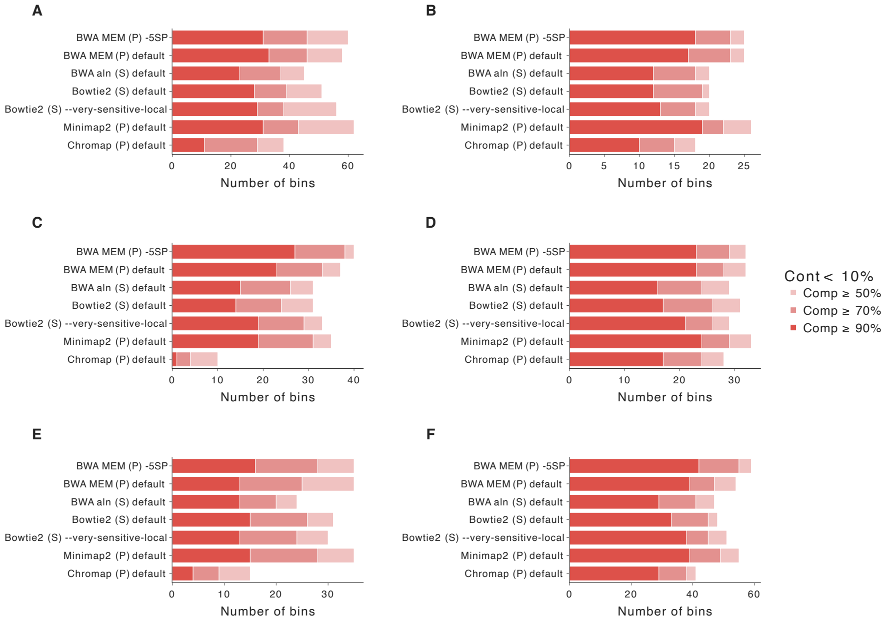


**Fig. 11 Binning results for human gut datasets.** The legend abbreviations are: "Cont" for Contamination and "Comp" for Completeness. Panels **A-F** show the number of bins with contamination $< 10\%$ and completeness thresholds of $\geq$ 50%, 70%, and 90% for the human gut datasets SPMP18 (**A**), SPMP19 (**B**), SPMP20 (**C**), SPMP21 (**D**), SPMP38 (**E**), and SPMP40 (**F**).

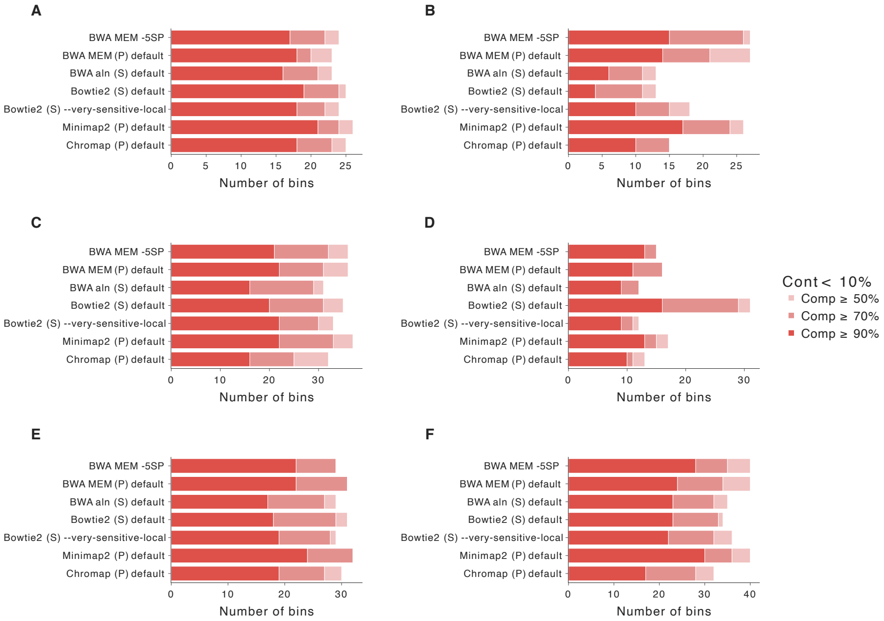


**Fig. 12 Binning results for human gut datasets.** The legend abbreviations are: "Cont" for Contamination and "Comp" for Completeness. Panels **A-F** show the number of bins with contamination $< 10\%$ and completeness thresholds of $\geq 50\%$, 70%, and 90% for the human gut datasets SPMP42 (**A**), SPMP50 (**B**), SPMP53 (**C**), SPMP58 (**D**), SPMP59 (**E**), and SPMP60 (**F**).

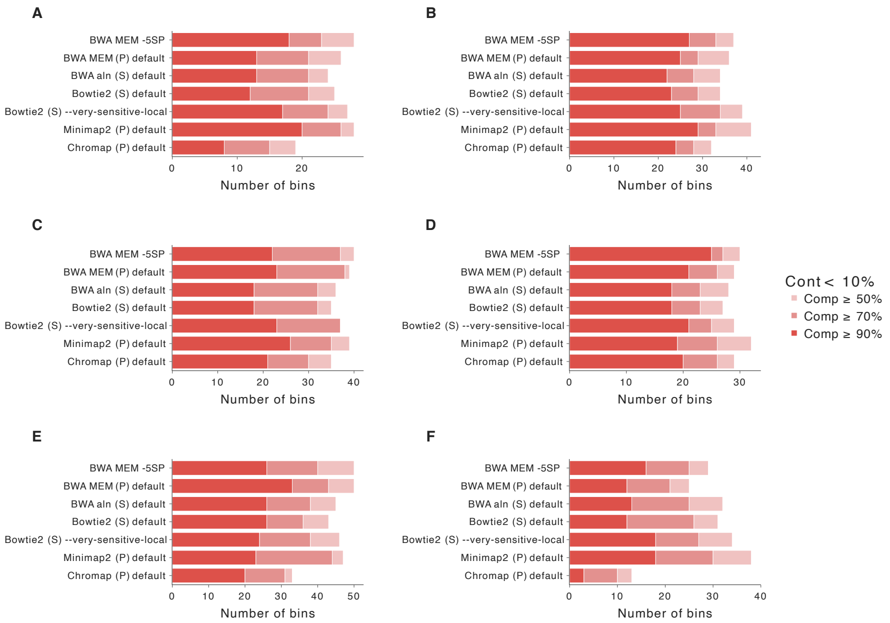


**Fig. 13 Binning results for human gut datasets.** The legend abbreviations are: "Cont" for Contamination and "Comp" for Completeness. Panels **A-F** show the number of bins with contamination $< 10\%$ and completeness thresholds of $\geq 50\%$, 70%, and 90% for the human gut datasets SPMP64 (**A**), SPMP67 (**B**), SPMP71 (**C**), SPMP73 (**D**), SPMP74 (**E**), and SPMP84 (**F**).

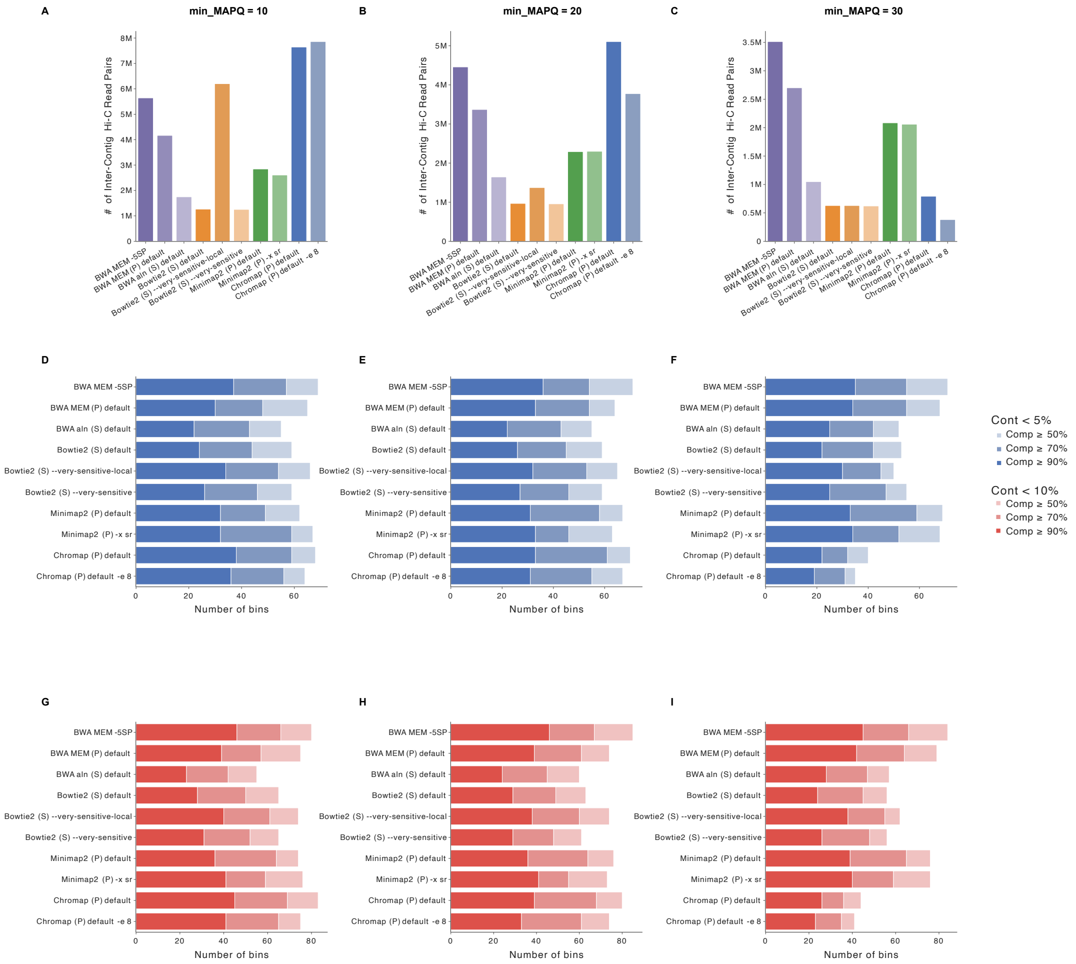


**Fig. 14 Downstream results across different MAPQ thresholds for bovine dataset.** "Cont" denotes contamination and "Comp" denotes completeness. Panels **A-C** show the number inter-contig Hi-C read pairs at different minimum MAPQ threshold. Panels **E-F** show the number of bins with contamination $< 5\%$ and completeness thresholds of $\geq 50\%$, 70%, and 90% across different MAPQ thresholds. Panels **G-I** show the number of bins with contamination $< 10\%$ and completeness thresholds of $\geq 50\%$, 70%, and 90% across different MAPQ thresholds.

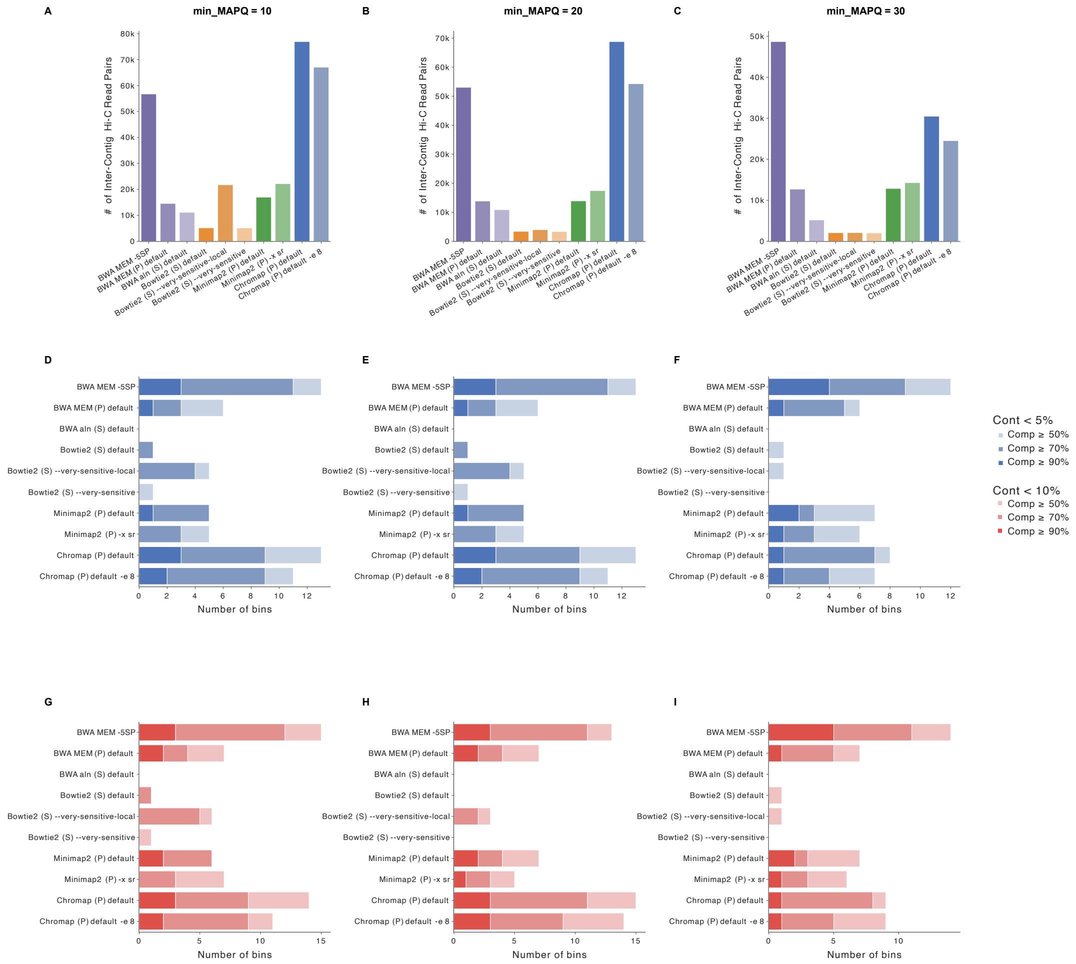


**Fig. 15 Downstream results across different MAPQ thresholds for human gut SPMP11 dataset.** "Cont" denotes contamination and "Comp" denotes completeness. Panels **A-C** show the number inter-contig Hi-C read pairs at different minimum MAPQ threshold. Panels **E-F** show the number of bins with contamination $< 5\%$ and completeness thresholds of $\geq 50\%$, $70\%$, and $90\%$ across different MAPQ thresholds. Panels **G-I** show the number of bins with contamination $< 10\%$ and completeness thresholds of $\geq 50\%$, $70\%$, and $90\%$ across different MAPQ thresholds.

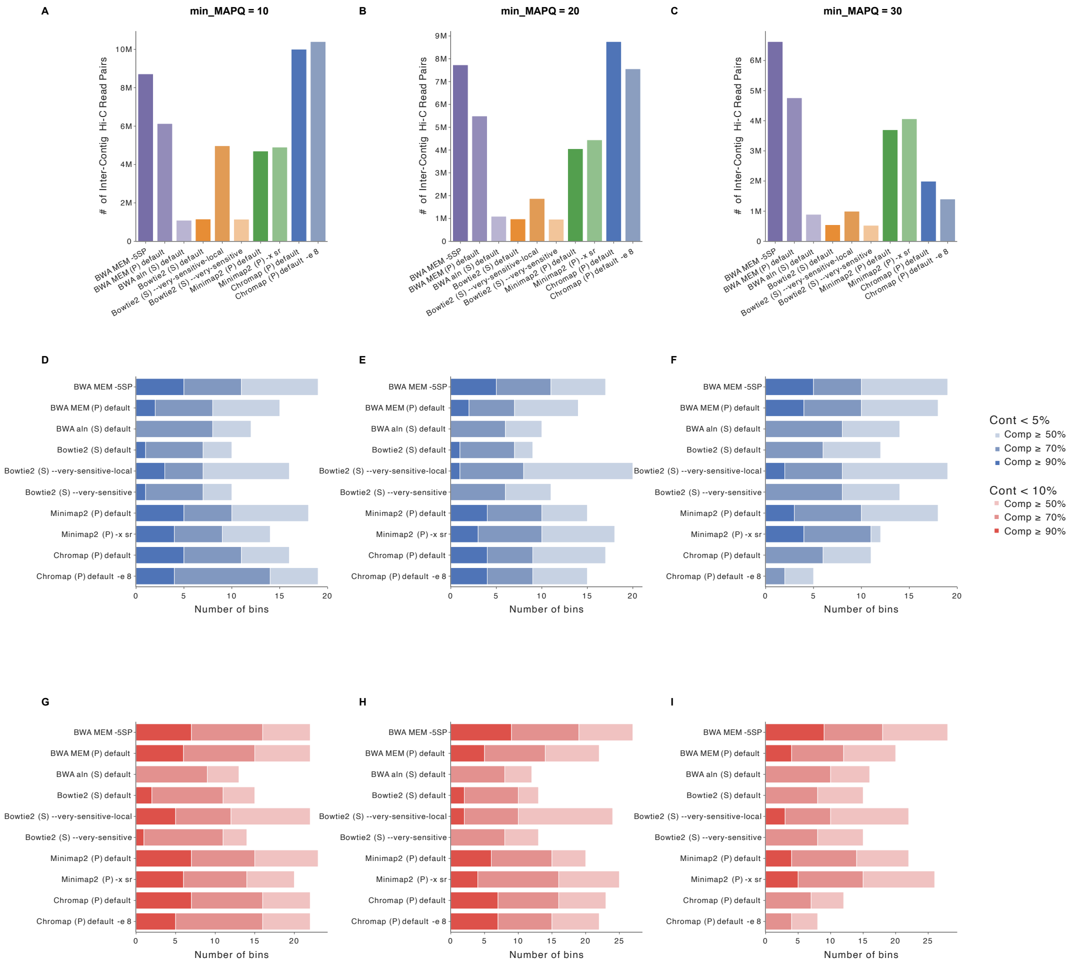

**Fig. 16 Downstream results across different MAPQ thresholds for hydrothermal mats M8 dataset.** "Cont" denotes contamination and "Comp" denotes completeness. Panels **A-C** show the number inter-contig Hi-C read pairs at different minimum MAPQ threshold. Panels **E-F** show the number of bins with contamination $< 5\%$ and completeness thresholds of $\geq 50\%$, 70%, and 90% across different MAPQ thresholds. Panels **G-I** show the number of bins with contamination $< 10\%$ and completeness thresholds of $\geq 50\%$, 70%, and 90% across different MAPQ thresholds.

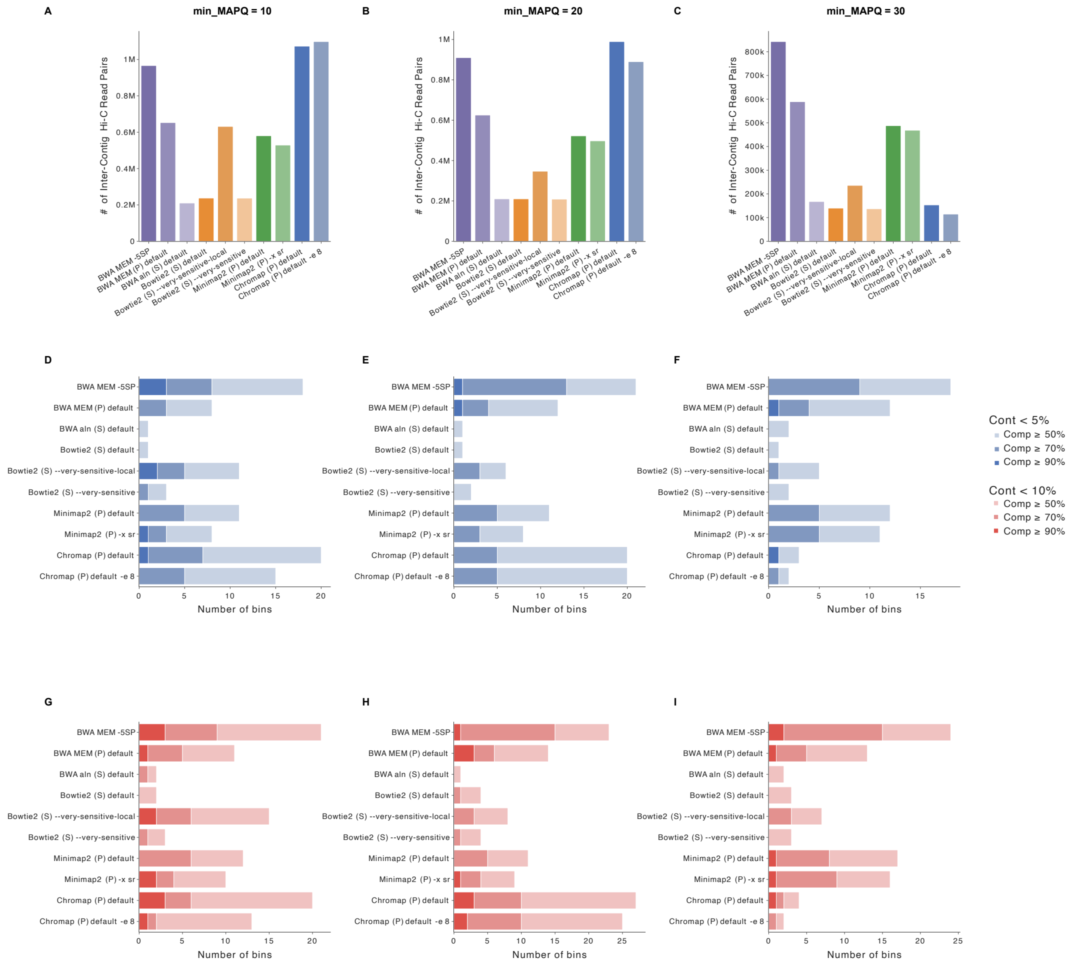


**Fig. 17 Downstream results across different MAPQ thresholds for pigs cv_7_1 dataset.** ”Cont” denotes contamination and ”Comp” denotes completeness. Panels **A-C** show the number inter-contig Hi-C read pairs at different minimum MAPQ threshold. Panels **E-F** show the number of bins with contamination $< 5\%$ and completeness thresholds of $\geq 50\%$, 70%, and 90% across different MAPQ thresholds. Panels **G-I** show the number of bins with contamination $< 10\%$ and completeness thresholds of $\geq 50\%$, 70%, and 90% across different MAPQ thresholds.

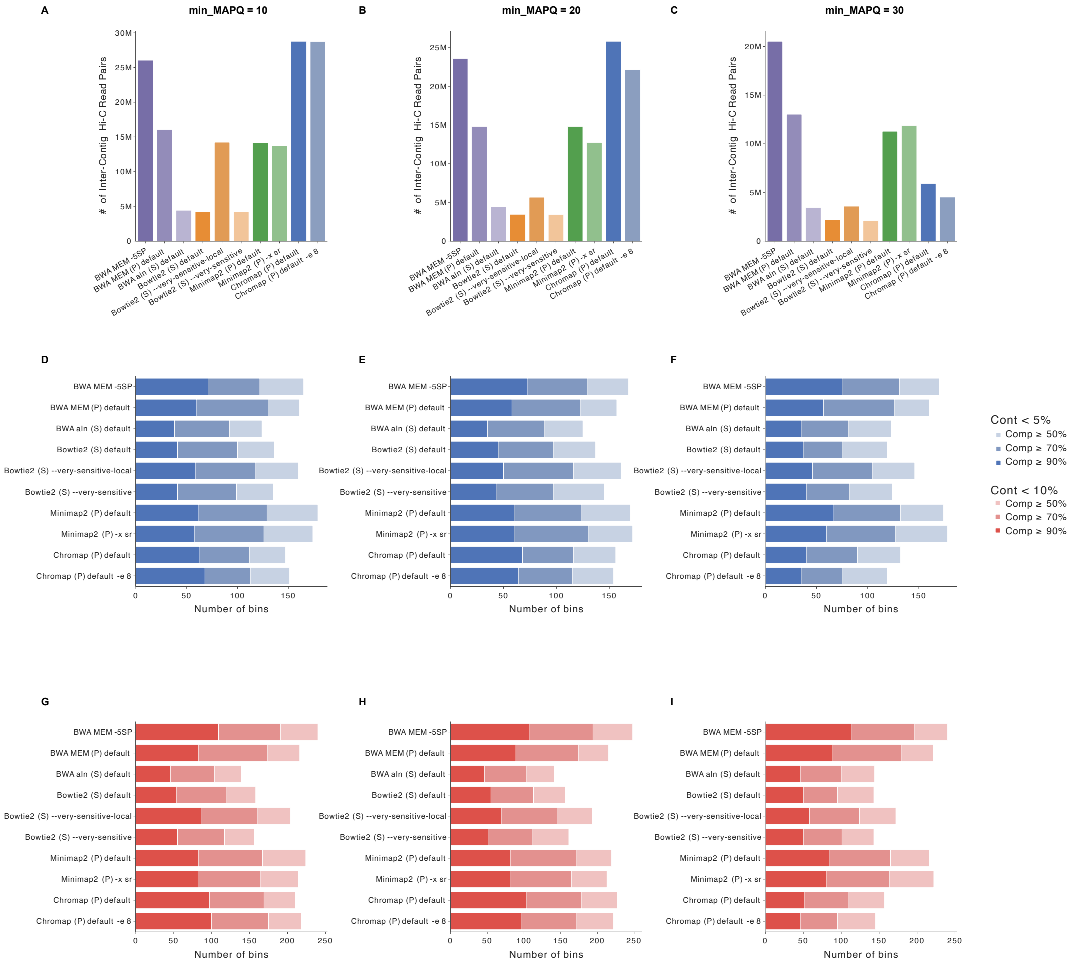


**Fig. 18 Downstream results across different MAPQ thresholds for wastewater dataset.** "Cont" denotes contamination and "Comp" denotes completeness. Panels **A-C** show the number inter-contig Hi-C read pairs at different minimum MAPQ threshold. Panels **E-F** show the number of bins with contamination $< 5\%$ and completeness thresholds of $\geq 50\%$, 70%, and 90% across different MAPQ thresholds. Panels **G-I** show the number of bins with contamination $< 10\%$ and completeness thresholds of $\geq 50\%$, 70%, and 90% across different MAPQ thresholds.

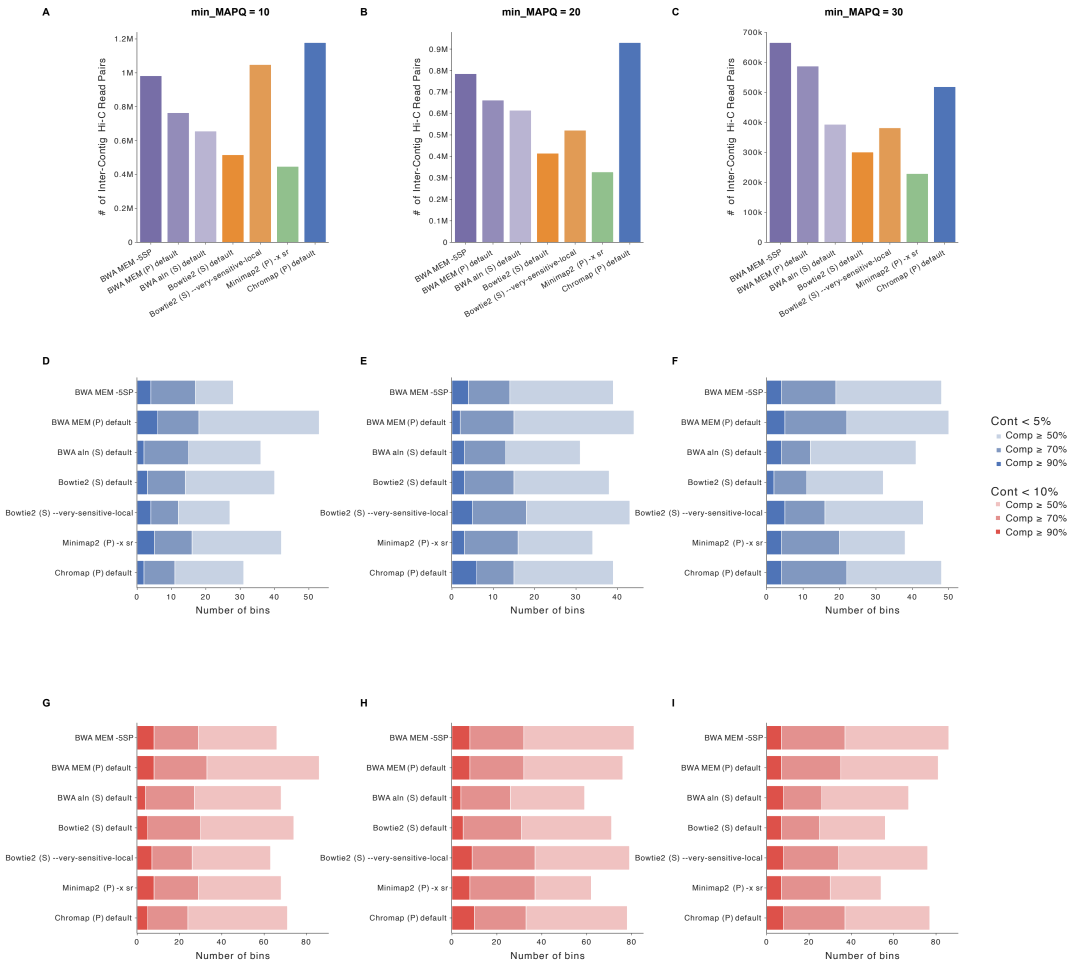


**Fig. 19 Downstream results across different MAPQ thresholds for cow rumen dataset.** "Cont" denotes contamination and "Comp" denotes completeness. Panels **A-C** show the number inter-contig Hi-C read pairs at different minimum MAPQ threshold. Panels **E-F** show the number of bins with contamination $< 5\%$ and completeness thresholds of $\geq 50\%$, 70%, and 90% across different MAPQ thresholds. Panels **G-I** show the number of bins with contamination $< 10\%$ and completeness thresholds of $\geq 50\%$, 70%, and 90% across different MAPQ thresholds.

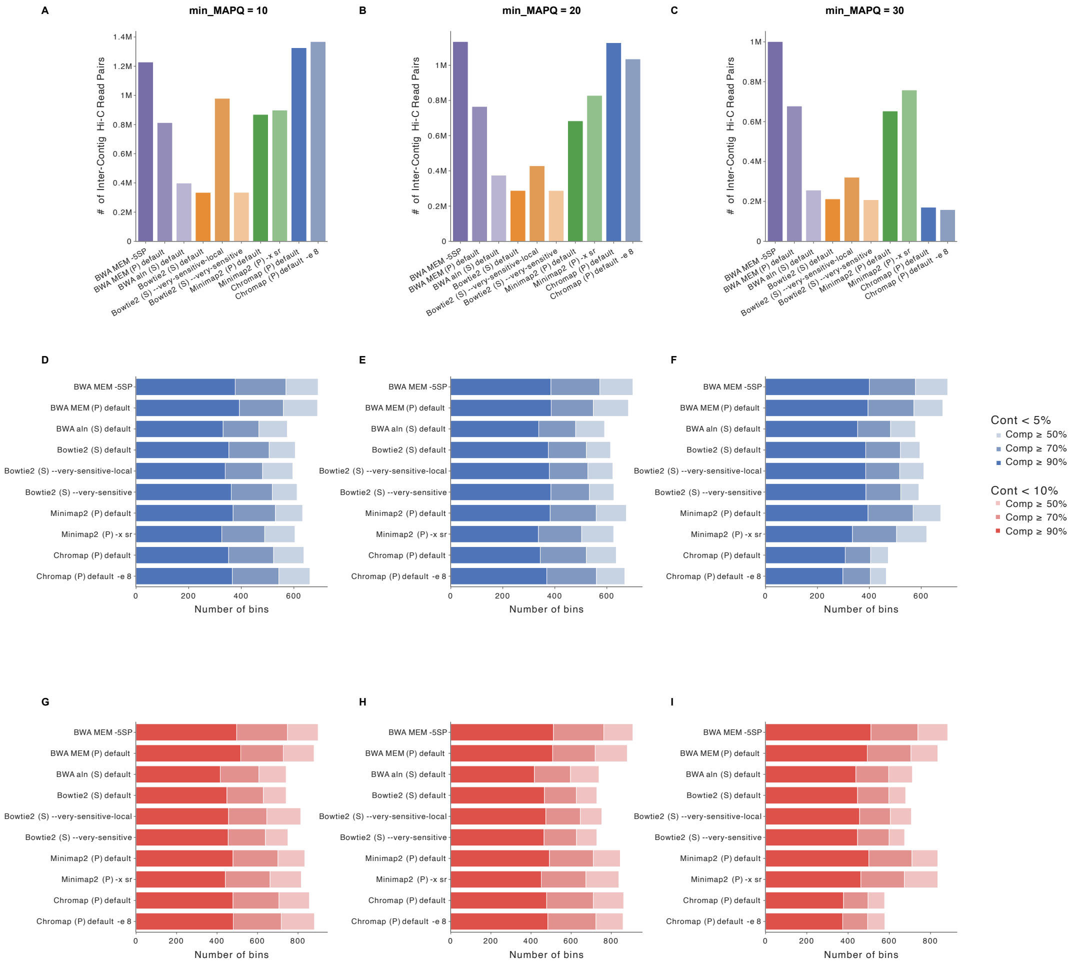


**Fig. 20 Downstream results across different MAPQ thresholds for sheep gut dataset.** "Cont" denotes contamination and "Comp" denotes completeness. Panels **A-C** show the number inter-contig Hi-C read pairs at different minimum MAPQ threshold. Panels **E-F** show the number of bins with contamination $< 5\%$ and completeness thresholds of $\geq 50\%$, 70%, and 90% across different MAPQ thresholds. Panels **G-I** show the number of bins with contamination $< 10\%$ and completeness thresholds of $\geq 50\%$, 70%, and 90% across different MAPQ thresholds.

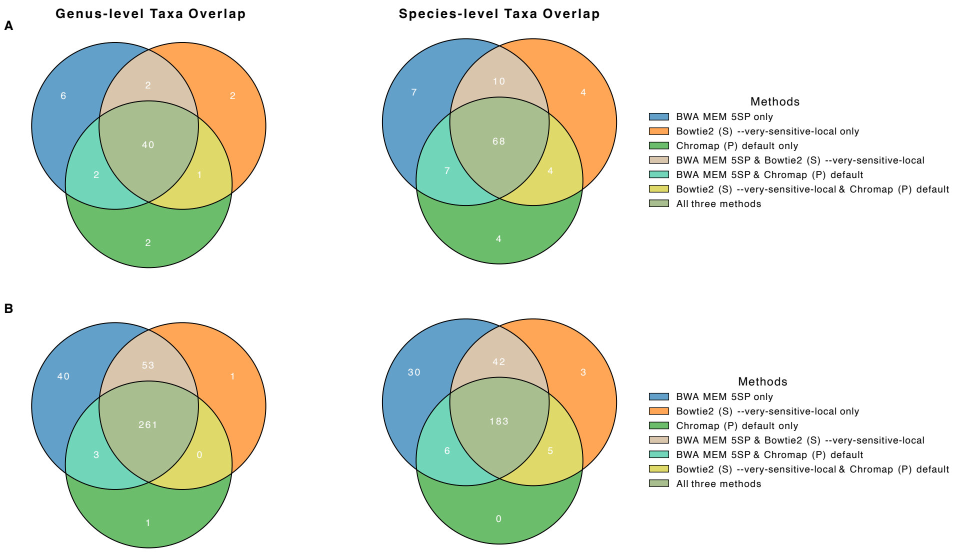


**Fig. 21 GTDB-Tk annotation results at the genus and species level.** Panels **A** shows the number of unique genus and species identified for the cow rumen dataset. **B** for the sheep gut datastet.

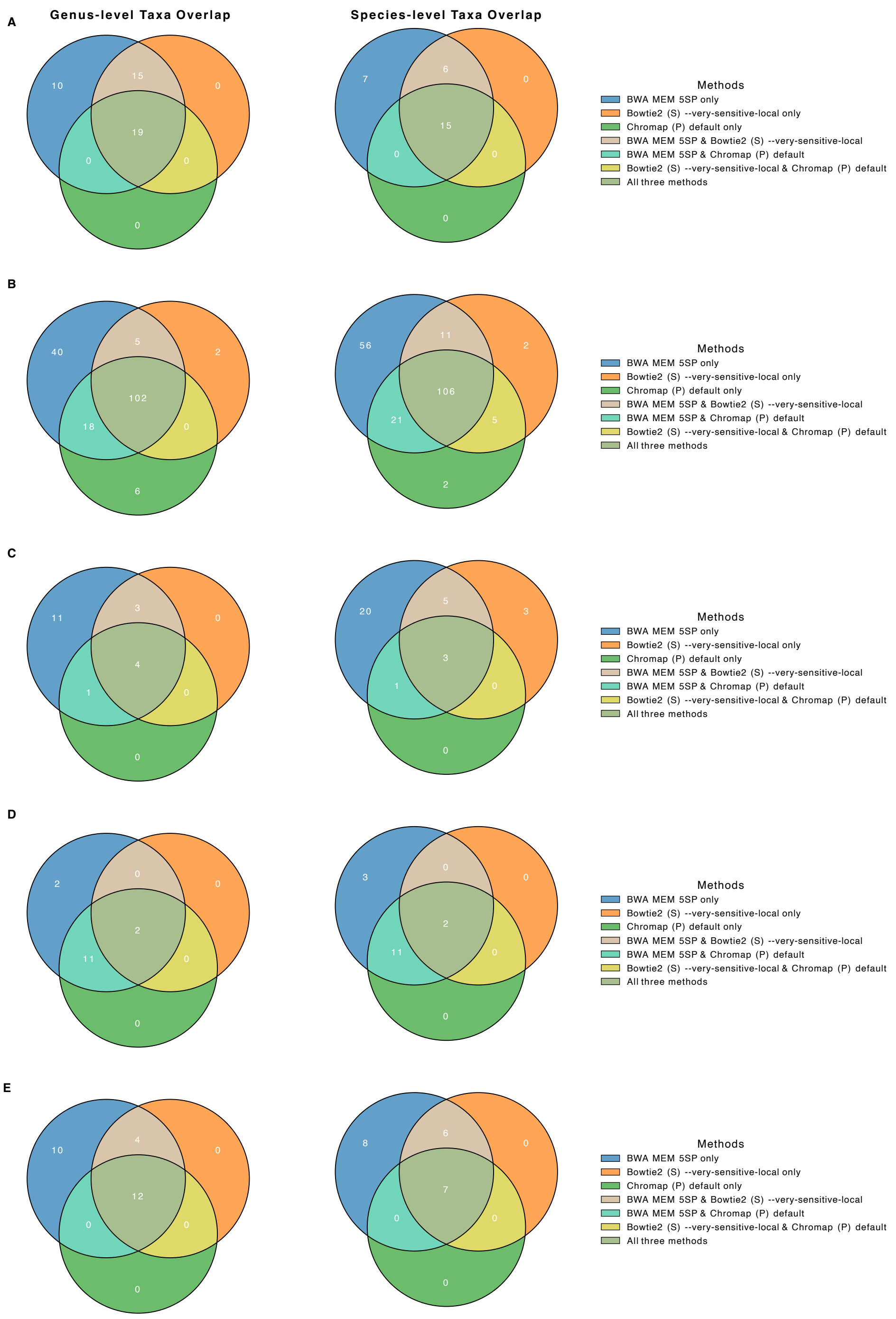


**Fig. 22 GTDB-Tk annotation results at the genus and species level.** Panels **A** shows the number of unique genus and species identified for the bovine dataset. **B** for the waste water datastet. **C** for the pig cv_7_1 datastet. **D** for the human gut SPMP11 datastet. **E** for the hydrothermal mats M8 datastet.
